# Negative impact of mild arid conditions in natural rodent populations revealed using markers of physiological condition in natura

**DOI:** 10.1101/2024.03.11.583554

**Authors:** Hamilcar Keilani, Nico Avenant, Pierre Caminade, Neville Pillay, Guila Ganem

## Abstract

1. Understanding how organisms respond to seasonal variations in their environment can be a window to their potential adaptability, a classical problem in evolutionary ecology. In the context of climate change, inducing increased aridity and disruption of seasonality, it is crucial to study the extent and limits of species responsiveness.
2. Here, the physiological response to food and water shortage during seasonally dry conditions were investigated. We studied populations of two rodent species of the genus *Rhabdomys*, one arid and one mesic, in a semi-arid zone where their range overlap in South Africa. We measured blood concentrations of markers of kidney and liver function, as well as body condition, at the onset and the end of the dry season.
3. We found similar shifts in blood metabolite levels, in the semi-arid populations of the two species, indicating malnutrition consistent with the observed degradation of habitat quality between the start and the end of the dry season. Furthermore, regardless of the period, differences between the two species in blood metabolite concentrations (e.g. amylase, sodium, alkaline phosphatase) were observed, suggesting contrasting diets and water conservation abilities.
4. Overall, we show that, as seasonal dry conditions worsen, organisms are increasingly affected by reduced food availability, and local adaptation to arid conditions may provide the arid species with an advantage to cope with semi-arid conditions. Our study suggests that even mild arid conditions could have a negative impact and questions resilience of animals to harsher arid conditions.

## Introduction

Organisms are constantly exposed to a wide range of environmental variations, including short-term changes during their lifetime and longer-term changes across generations (Lopez-Maury et al., 2008). In the coming decades, extreme and more frequent droughts are predicted by most projections, resulting from changes in precipitation patterns, increased temperature, and evaporation (Naumann et al., 2018). These changes will induce great challenges to organisms. Under such circumstances, divergent selection can lead to the evolution of local adaptations where, in a given environment, resident phenotypes outperform nonresident ones (Kawecki & Ebert, 2004).

As a result of climate change, organisms can be faced with warmer and drier environmental conditions (Parmesan et al., 2000), likely to impact their capacity to maintain homeostasis (i.e. the state of steady internal conditions allowing optimal body functioning) (Davies, 2016; Fuller et al., 2016). In dry environments, food and water can be scarce, generating strong selection on physiological attributes maximizing energy and water availability for body function. Habitat generalists and specialists’ taxa may respond differently to environmental changes. Desert specialists exhibit specific physiological and/or behavioral traits, resulting from genetic adaptations to the environment, allowing them to continuously endure or evade periods of resource restriction (Rocha et al., 2021). In contrast, generalist species may seek to escape the effects of aridity through temporary avoidance (Abraham et al., 2019) or plastic adjustments during unfavorable seasons (Kobbe et al., 2011). Compared to mesic species, arid species have lower basal metabolic rates and evaporative water loss (Muñoz-Garcia et al., 2022), adopt a more flexible diet (Tshikae et al., 2013), and cope better with prolonged droughts (Boyers et al., 2021).

Local ecological adaptation can be described in several different and complementary ways, such as through its genetic basis, transcriptomic, behavioral, or physiological aspects. Adaptive evolution *sensu stricto* occurs when the genetic constitution of a population changes because of natural selection (Merilä & Hendry, 2013). In contrast, local adaptation *sensu lato*, also considers concepts such as phenotypic plasticity (the ability of a genotype to produce distinct phenotypes when exposed to different environments throughout its ontogeny; Pigliucci, 2005). It is usually assessed using comparative population or species fitness estimates under novel environmental conditions (Webster & Reusch, 2017). The response to local environmental conditions is a major factor in the generation and maintenance of biodiversity, impacting population dynamics, biogeographical ranges, and species interactions (Blanquart et al., 2013; Post et al., 2009). Specifically, physiological limits can drive and be driven by evolution, shape species distributions and niches, and define species response capacities to future climate change, directly impacting risks of extinction (Somero, 2012). For example, latitudinal niches are associated with adaptive variation in thermal limits in marine invertebrates (Somero, 2010).

Studies of different physiological systems, such as the cardiovascular (Seebacher et al., 2005, Berkel & Cacan, 2022) or digestive systems (Naya et al., 2011), in response to different abiotic conditions, can improve our understanding of the causal mechanisms of responses of organisms to climate variation (Seebacher & Franklin, 2012). Biotic factors, such as sex, age, diet, strain, or breeding status also influence individual responses (McClure, 1999). With a focus on influence of air temperature, water availability and energy resources on an organism’s physiological state, research in physiological ecology has addressed how variation in physiology evolves and is maintained in relation to environmental conditions (Feder & Block, 1991).

Analysis of blood metabolites has successfully been used to investigate kidney and liver physiology of wild populations (e.g. Al-Eissa et al., 2012). The kidney and the liver both contribute to homeostasis in periods of food and water scarcity; indeed, the kidney plays a crucial role in maintaining osmotic balance and the liver is involved in dietary metabolism. Water loss can also mechanically induce high concentrations of some blood markers and impact organism’s capacity to evacuate products of metabolic waste, such as urea, from the blood (Ostrowski et al., 2006). Hence, dry conditions can put a strain on these organs and be detrimental to survival and reproduction.

In the Succulent Karoo of South Africa, where the average daily precipitation during the dry season is 0.26 mm *versus* 0.78 mm during the wet season, the probability of survival of adult *Rhabdomys pumilio,* a species of African striped mouse, were related to their physiological response at the start of the dry season (Schoepf et al., 2017a). Schoepf and collaborators found higher serum concentrations of albumin, glucose, potassium, and lower concentrations of globulin, in animals that survived compared to those that did not survive the dry season. At the peak of the dry season, individuals had lower concentrations of glucose and phosphorus and higher concentrations in globulin and urea nitrogen if they survived the dry season. These results strongly suggest that such markers could be good indicators of individual fitness.

In our study, potential impact of increased aridity was addressed by investigating the physiological consequences of seasonal variation in dry conditions, in two species of the diurnal African striped mouse genus *Rhabdomys*. The two study species, *R.bechuanae* and *R.dilectus dilectus*, have different environmental niches (du Toit et al. 2012, Meynard et al. 2012). Indeed, throughout most of its distribution, *R.d.dilectus* occurs in mesic habitats with dense ground vegetation cover and nests in dense grass, while *R.bechuanae* is found in semi-arid and arid habitats, thriving predominantly in sparsely vegetated areas and nesting in bushes (Dufour et al., 2015; Dufour et al., 2019). Moreover, differences in morphology and behavior suggest adaptation to dry conditions in *R.bechuanae* (Ganem et al., 2020; Dufour et al., 2019). Here we address both physiological responses to seasonal dry conditions and divergence between populations of the two species in the same bioclimatic region.

Indeed, at the edge of their distributions in central South Africa, the two species inhabit a semi-arid region within which they occur either as parapatric or sympatric populations. This semi-arid zone, the expansion and contraction of which depends on land use and precipitation (Lian et al., 2021), has experienced a recent trend towards desiccation (Jury, 2021). Such drying conditions could generate additional selective pressures to those already experienced by the semi-arid populations of the two species. Short-term climatic patterns can also impact resource availability in this region. Indeed, during La Niña years (like during this study), as part of the El Niño Southern Oscillation phenomenon, there is a general association between regional wetness and sea surface temperatures in the neighbouring Atlantic and Indian Oceans, leading to wetter episodes throughout Southern Africa (Nicholson & Selato, 2000).

While many studies have harnessed physiology, genomics, and transcriptomics to highlight potential adaptive interspecific or interpopulational contrasts, studies under common natural environmental conditions are lacking (Rocha et al., 2021). These approaches provide a mean to disentangle the roles of interpopulational or interspecific variation from extrinsic environmental factors in shaping phenotypic variation, taking advantage of the natural conditions experienced in the field, including their complexity, with little human involvement. This study took place in a natural semi-arid environment and asked how seasonal variation in dry conditions influenced the physiological responses of *R.bechuanae* and *R.d.dilectus*. We compared body condition, blood concentrations of markers of kidney and liver functions and habitat characteristics of parapatric populations of the two species at the start *versus* the end of the dry season. First, we expected that the habitat available for the mice would be drier at the end compared to the onset of the dry season, inducing a reduction of food and water, impairing liver and kidney function, and impacting body condition. Second, assuming local adaptive plasticity, we hypothesized that both species would be able to adjust their physiological responses to seasonal changes in dry conditions. Third, we expected that *R.bechuanae*, having evolved in arid environments, would perform better in semi-arid conditions and be closer to the expected local optimal response to increased dry conditions than *R.d.dilectus* (having evolved in mesic areas), as found for *Mus musculus* (Bittner et al., 2021). Alternatively, if both species evolved specific adaptations to the semi-arid environment, we expected no species differences (**Table 1**).

**Table 1:** Summary of main ecological characteristics of R.bechuanae and R.d.dilectus and of our predictions on their physiological response during the dry season.

| ECOLOGICAL CHARACTERISTICS | *R.BECHUANA*E | *R.D.DILECTUS* |
| --- | --- | --- |
| **ENVIRONMENTAL NICHE** | Arid to semi-arid<br>(Ganem et al., 2020) | Mesic to semi-arid (Ganem et al., 2020) |
| **PREFERRED HABITAT** | Bushy patches<br>(Dufour et al., 2015) | Continuous cover / grassland<br>(Dufour et al., 2015) |
| **SOCIAL STRUCTURE** | Group-living<br>(Dufour et al., 2019) | Solitary / opportunistic group-living<br>(Dufour et al., 2019) |
| **SPECIES-SPECIFIC RESPONSE TO SEASONAL VARIATIONS OF DRY CONDITIONS: PREDICTIONS** |  |  |
| **BODY CONDITION** | ↘↘ or ↘* | ↘↘ |
| **NUTRITIONAL METABOLISM** | ↘↘ or ↘* | ↘↘ |
| **LIVER ACTIVITY** | ↘↘ or ↘* | ↘↘ |
| **WATER BALANCE** | ↘↘ or ↘* | ↘↘ |
| **SENSITIVITY TO INFECTION** | ↗↗ or ↗* | ↗↗ |
| **SPECIES DIFFERENCES: PREDICTIONS** |  |  |
| **BODY CONDITION** | + | - |
| **LIVER ACTIVITY** | + | - |
| **WATER BALANCE** | + | - |
| **SENSITIVITY TO INFECTION** | - | + |

\*Adaptation to harsher arid conditions could have enabled *R.bechuanae* to withstand growing aridity without suffering as many adverse effects as *R.d.dilectus* in the semi-arid zone. Alternatively, semi-arid populations of both species could both have developed local adaptation to dry conditions.

## Materials and Methods

### Ethics statement

Permits to sample and handle animals in the field were obtained from the DESTEA of the Free State, the North West DEDECT and Northern Cape DENC (respectively, n°202110000007994, NW 38956/04/2022, FAUNA 0389/2022). Animal handling was performed under ethical clearance from the Languedoc-Roussillon ethical committee for animal experimentation (n°2022041512256467 v5).

### Choice of sampling periods and sites

We selected six study sites based on their geographical position and their aridity index (AI). The AI calculation was based on Thornthwaite method (Thornthwaite, 1948; see **Supplementary Material** for full formula) using rainfall and temperature data collected from 2010-2021 (South African Weather Service).

We selected parapatric populations of the two species in a semi-arid region of South Africa (0.2 < AI < 0.5). In this region, a hot and wet season occurs from roughly December to February, and a cool dry season from May to September. We sampled six sites at the onset (May 2022) of the dry season: Barberspan Bird Sanctuary, Benfontein Nature Reserve, Bloemhof Dam Nature Reserve, Gariep Dam Nature Reserve, Kalkfontein Dam Nature Reserve and Wolwespruit Nature Reserve (**Figures 1-3, Table 2**). We then resampled four of these sites at the end of the dry season (September 2022). Two sites (Barberspan and Benfontein) were excluded from the September sampling due to extreme weather conditions.

**Figure 1:**
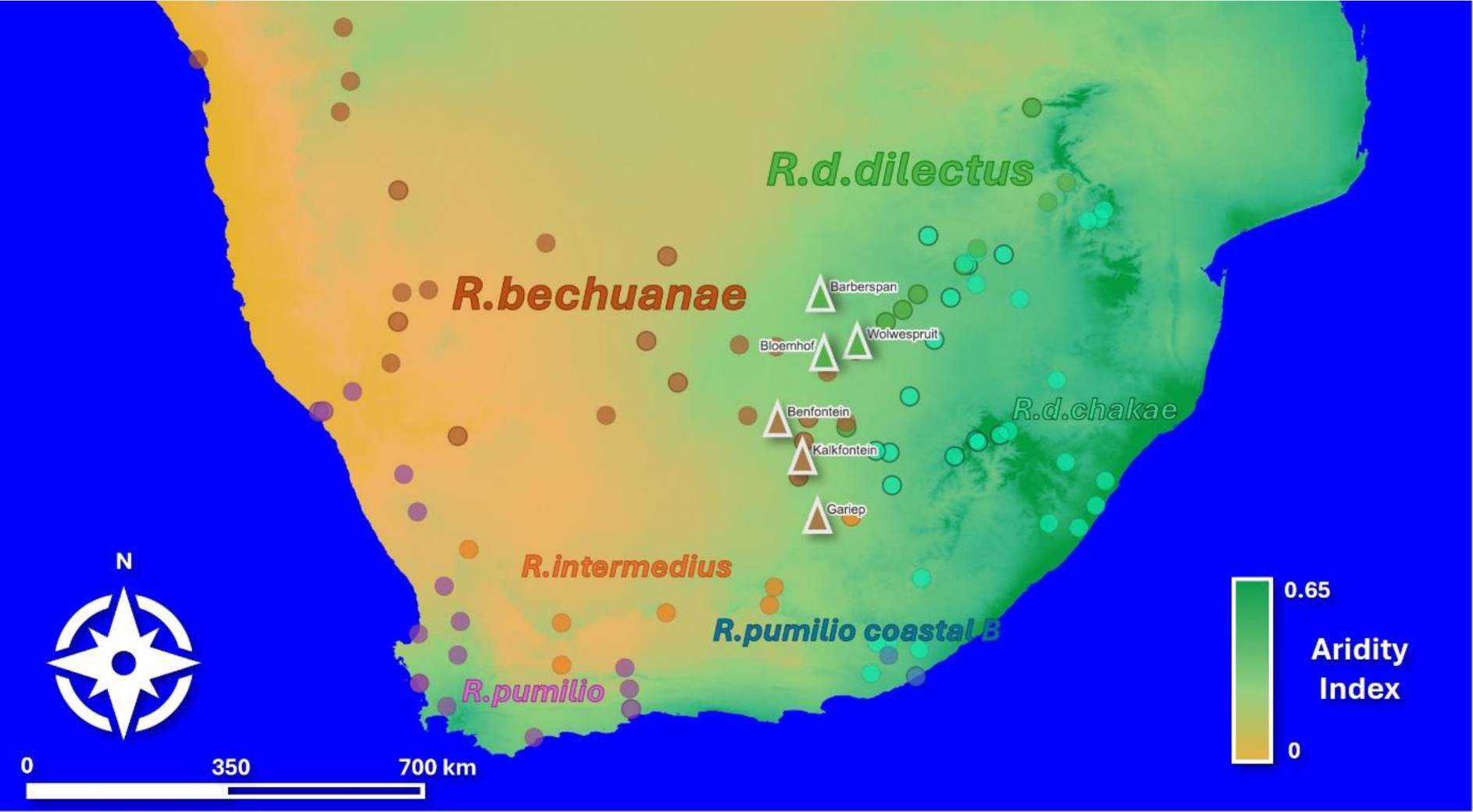
Map of known occurrences of 6 main clades of Rhabdomys (translucent dots) and locations sampled in this study (full colour dots) in southern Africa. Based on published data and unpublished data (origin details are available in doi: 0d1b3414-7e2a-11ea-a38d-00163e26bfb0). Base map: World Topographic Map Esri Standard, Aridity Index layer was computed from a 0.5° global grid, using data from Version 3 of the Global Aridity Index and Potential Evapotranspiration Database (Zomer et al.,2022).

**Table 2:** Geographical position and Aridity Index of sampled sites for each species.

| SPECIES | SAMPLED SITES | GEOGRAPHICAL COORDINATES | ARIDITY INDEX<sup>*</sup> |
| --- | --- | --- | --- |
| ***R.bechuanae*** | Gariep Dam Nature Reserve | E25.55537; S30.57589 | 0.42 ± 0.07 |
|  | Kalkfontein Dam Nature Reserve | E25.28031; S29.52058 | 0.54 ± 0.07 |
|  | Benfontein Nature Reserve** | E24.81824; S28.82747 | 0.39 ± 0.07 |
| ***R.d.dilectus*** | Bloemhof Dam Nature Reserve | E25.65144; S27.66063 | 0.35 ± 0.06 |
|  | Barberspan Bird Sanctuary** | E25.58354; S26.56538 | 0.39 ± 0.05 |
|  | Wolwespruit Nature Reserve | E26.26326; S27.41418 | 0.53 ± 0.10 |

\*Average Aridity index (see **Supplementary material** for formula) ± Standard Error of the Mean, based on climate data collected from 2010 to 2021 (South African Weather Service)
\*\* Sampled only in May 2022 (dry season onset).

### Capture and field protocol

We used small mammal PVC traps (LxHxW= 29.6×7.5×7.5 cm), baited with a mixture of oats, peanut butter, salt and sunflower seeds, and added a piece of cotton wool. They were placed approximately every 15 m along roughly 150 to 300 m transects. Number and length of transects varied with habitat and terrain conditions. Overall, trapping effort was 9688 trap nights (details in **Supplementary Tables 1&2**). Since *Rhabdomys* is mostly diurnal, the traps were checked twice a day (8 AM and 3 PM). On the field, all trapped striped mice were measured (body length and mass), and their sex and breeding status (breeding or non-breeding) assessed based on external morphological features; individuals considered as breeders presented either signs of lactation, a perforated vagina, a vaginal plug, or scrotal testes. Additionally, a 0.5 mm piece of tail was collected and kept in 98 % ethanol for species identification. All striped mice were marked with a unique ear-tag before their release at the trapping location unless they were kept for the physiological study (i.e. kept in their trap with cotton wool and food).

Out of 903 trapped small mammals, 694 were *Rhabdomys*. For the physiological study, we selected only adults, avoiding related individuals as much as possible, by selecting mice trapped at least 100 m apart, except for breeding pairs (a male and a female) that could be trapped in the same nest. This distance was based on data obtained for *R. pumilio* indicating minimal relatedness (R=0 between males and R=0.06 between females) at this distance (Solmsen et al. 2012). In each site, trapping lasted 3 to 6 days.

### Habitat

#### Vegetation composition

Earlier studies have shown inter-species differences in vegetation cover and structure requirements (Dufour et al., 2015). Since habitat characteristics can vary locally, we characterized the vegetation structure (i.e. grass *versus* woody vegetation) at a microhabitat scale (around the trap), and at a mouse home-range scale (Dufour et al., 2015). Around traps in which a *Rhabdomys* was trapped, the percentage of the surface composed of dry grass, green grass, dry bushes, green bushes, succulent plants, holes and uncovered surfaces was assessed within 2mx2m (4m^2^) and the general vegetation within 10×10m (100m^2^) quadrats centered on the trap position. For each 4m^2^ quadrat we recorded in detail the percentage and type of cover at the ground level within each of four 1×1m subunits (using a metal frame); the results obtained for the four subunits were then averaged. For the 100m^2^ quadrats the assessment was made following a visual inspection at eye-level height; the different cover percentages were estimated by the same observer. All together we characterized 236 quadrats of each type (100m^2^ and 4m^2^).

#### Normalised Differential Vegetation Index

NDVI, the satellite imagery-based index informative of ground vegetation greenness, was retrieved from the Copernicus Open Access Data Hub (Copernicus Sentinel-2 data, 2023, calculated from 10-m resolution bands). While the two above-mentioned quadrats were aimed at characterizing the structure/cover of the habitat, NDVI was used as an index of habitat quality. To characterize each site and sampling session, we used NDVI data available for the closest day to the beginning of a sampling session, which was expected to represent the conditions experienced by the mice at the time of capture.

### Blood extraction and assessment of blood metabolites concentrations

In this study, a total of 273 adult mice were euthanized by means of cervical dislocation on their day of capture, and a blood volume of around 100 µL was collected in lithium-heparinized tubes by cardiac puncture. The breeding status was confirmed for all individuals during dissection. The whole liver, the left kidney, and the skull of every individual were also collected for a complementary study. Because the two species could not be distinguished visually, the spleen was also harvested from each mouse and subsequently analyzed for post-hoc species identification using Cytochrome Oxidase I genotyping (as described in Ganem et al. 2020).

Levels of albumin, alkaline phosphatase, alanine aminotransferase, amylase, total bilirubin, blood urea nitrogen, total protein, globulin, glucose, calcium, phosphorus, potassium, and sodium were successfully measured in the whole blood of 257 individuals immediately after blood collection (Abaxis VETSCAN V2 technology with the Comprehensive diagnostic profile cartridge), following the manufacturer’s instructions. The coefficient of variation (CV%) of measured levels of these markers was determined from data obtained by analyzing a given blood sample twice. Samples from six adult striped mice (5 *R.bechuanae*, 1 *R.d.dilectus*) were used for these calculations. For these six samples, the calculated CV equaled 5.6 ± 1.9 %, which is consistent with the intra-test CV% provided in the manufacturer’s instruction manual for other mammal species (**Supplementary Table 3**).

### Data preparation and analysis

All statistical tests were carried out using the R software (version 4.2.1).

#### Habitat

##### Vegetation composition

Because the distribution of the variables describing the cover and vegetation type did not meet the conditions necessary for further multivariate model based analyses, we performed a Principal Component Analysis (package FactoMineR version 2.9) including the 7 variables measured within the 4m^2^ quadrats and within the 100m^2^ quadrats aiming to transform the variables in a way that would take into account each vegetation structure parameter’s contribution to the overall variance. In each case, over 80 % of the variance was explained by the first five Principal Components (**Supplementary Tables 4 & 5**). The distribution of the coordinates of all traps on the first 5 PCs complied to the statistical analysis constraints and were used as response variables of a *PERMANOVA* (package *vegan*, version 2.6-4). *PERMANOVA* is a multivariate statistical inference tool using permutational algorithms (Anderson, 2001). This equivalent to *MANOVA* operates in a distribution-free setting and is robust to non-normality of residuals as well as dispersion heterogeneity, even in slightly unbalanced designs (Anderson, 2013). We tested differences between sites nested within sessions (start and end of dry season), with 10 000 permutations, to assess whether the sites had similar enough habitat characteristics to be considered as suitable replicates in other analyses.

##### NDVI

NDVI was averaged over a 60 m radius circular buffer centered at each trap and then compared across sessions and sites using a GLM Analysis of Variance, with the site nested within session. To meet statistical test prerequisites, the response variable was log-transformed; the analysis was followed by Tukey multiple comparisons of means post-hoc tests.

###### Age classes

Among trapped *Rhabdomys*, only adults were selected for this study based on their size, general appearance (fur dullness, presence of scars), and/or breeding status. Body length of trapped individuals varied between 5 and 12 cm. The body length of adult individuals involved in the physiological study varied between 7.7 and 12 cm. To distinguish between older and younger adults, mass was plotted against body length (**Supplementary figure 6**) showing a curve defined by an allometric growth equation with a steeper slope at the highest and lowest body length values. Since bone growth can significantly alter the concentration of some physiological markers, and growth rate varies with age/body length, we considered 4 length/age classes among the adults (Class A: [7.7:9 cm[; Class B : [9:10 cm[; Class C : [10:11 cm[; Class D [11 cm:]).

###### Body condition

We calculated a scaled mass index of body condition using the method described in Peig and Green (2009; see **Supplementary Material** for full formula), to assess the state of each individual’s fat reserves (Schulte-Hostedde et al., 2005). To test whether breeding status, age, habitat quality, increased dryness, and/or interspecific differences influence body condition of studied individuals, linear mixed effect models were computed with body condition as a response variable, site as a random factor and session, species, sex, session*species*sex (including all two-way interactions), breeding status (nested within sex), age class and NDVI as explanatory variables. Assumptions of normality and homoscedasticity of residuals were checked using diagnostic plots (*graphics* package v.3.6.2.). A Tukey multiple comparisons of means post-hoc test was applied to determine which inter-level differences were driving the significance of each factorial effect having more than two modalities.

###### Physiology

We first identified twelve outliers that we removed from further analyses. These outliers were animals from which we collected smaller volume of blood samples than recommended for the Vetscan or were heavily hemolyzed and flagged as such in the Vetscan analysis output. In addition, as total protein values were used by the Vetscan analyzer to infer the Globulin (GLOB) fraction of blood protein (with GLOB=TP-ALB), we considered this marker as not informative and removed this variable from our analyses. Finally, for 199 out of 257 samples, blood concentrations of creatinine (CRE) were flagged as being below the instrument’s sensitivity threshold; we hence excluded CRE from subsequent analyses, as this imprecision would have hindered analytical performance.

Due to covariance in variables such as blood physiological markers, multivariate statistical techniques provide a unique insight into the main patterns and effects driving variation in the data. As with habitat, multivariate normality was violated in our physiological dataset, so permutational multivariate analysis of variance (*PERMANOVA*) was performed. 10 000 permutations were used, with 236 blood samples, 12 physiological parameters as response variables, site as a random factor, and session, species, sex, breeding status (nested within sex), body condition, age class, and NDVI as explanatory variables. A backwards stepwise model selection was performed to retain the most parsimonious model. We then sought to identify which of the 12 physiological markers drove significantly the between-group differences evidenced with the *PERMANOVA*. Because in the PERMANOVA package a dedicated function was not available, twelve post-hoc univariate linear mixed model (LMM) tests were performed (followed by backwards stepwise model selection), initially using the parameters of the most parsimonious *PERMANOVA* model. Assumptions of normality and homoscedasticity of residuals were checked using diagnostic plots. In the case of total bilirubin, a permutation test had to be performed instead of the *LMM*. Through a backwards stepwise model selection process. For all tests, the significance level (α) was set at 0.05.

## Results

### Variation in habitat characteristics between species and sessions

#### Vegetation composition

We compared the vegetation characteristics surrounding each successful trap with a *PERMANOVA*, using the first 5 Principal Component coordinates (**Supplementary Tables 4 & 5**) of each quadrat as response variable, and site and session as explanatory variables. We detected significant differences in overall habitat characteristics between sessions (4m^2^ and 100m^2^: p<0.001), and sites nested within sessions (4m^2^ and 100m^2^: p<0.001), indicating clear site differences and seasonal effects on habitat structure (**Supplementary Figures 4 & 5**). Although a permutational dispersion test indicated that the latter patterns could be attributable at least in part to differences in dispersion between sessions or sites (see **Supplementary Material : Tables 6&7; Figures 1,2,3&4**), some general trends could be inferred from the data (**Supplementary Tables 7 & 8**). Indeed, quadrats from *R.d.dilectus* sites (Barberspan, Bloemhof and Wolwespruit) tended to differ from *R.bechuanae* ones on Principal Components most characterised by variance in grass cover, such as PC1 (mean 4 m^2^ quadrat coordinates: *R.d.dilectus* −0.68 ± 0.07, *R.bechuanae* 0.79 ± 0.15; mean 100 m^2^ quadrat coordinates : *R.d.dilectus* −0.59 ± 0.10, *R.bechuanae* 0.64 ± 0.13) and PC2 (mean 4 m^2^ coordinates : *R.d.dilectus* 0.24 ± 0.09, *R.bechuanae* −0.29 ± 0.11; mean 100 m^2^ coordinates : *R.d.dilectus* −0.31 ± 0.10, *R.bechuanae* 0.34 ± 0.11), or bush cover, such as PC4 for 100 m^2^ quadrats (mean coordinates : *R.d.dilectus* −0.27 ± 0.10, *R.bechuanae* 0.16 ± 0.09). Differences between sessions could also be observed on Principal Components most characterised by variance in grass cover, such as PC1 at the 100m^2^ level (mean coordinates : onset −0.11 ± 0.09, end 0.84 ± 0.07) and PC2 (mean 4m^2^ coordinates: onset 0.27 ± 0.11, end −0.38 ± 0.04; mean 100 m^2^ coordinates: onset −0.60 ± 0.11, end 0.84 ± 0.07).

**Figure 2:**
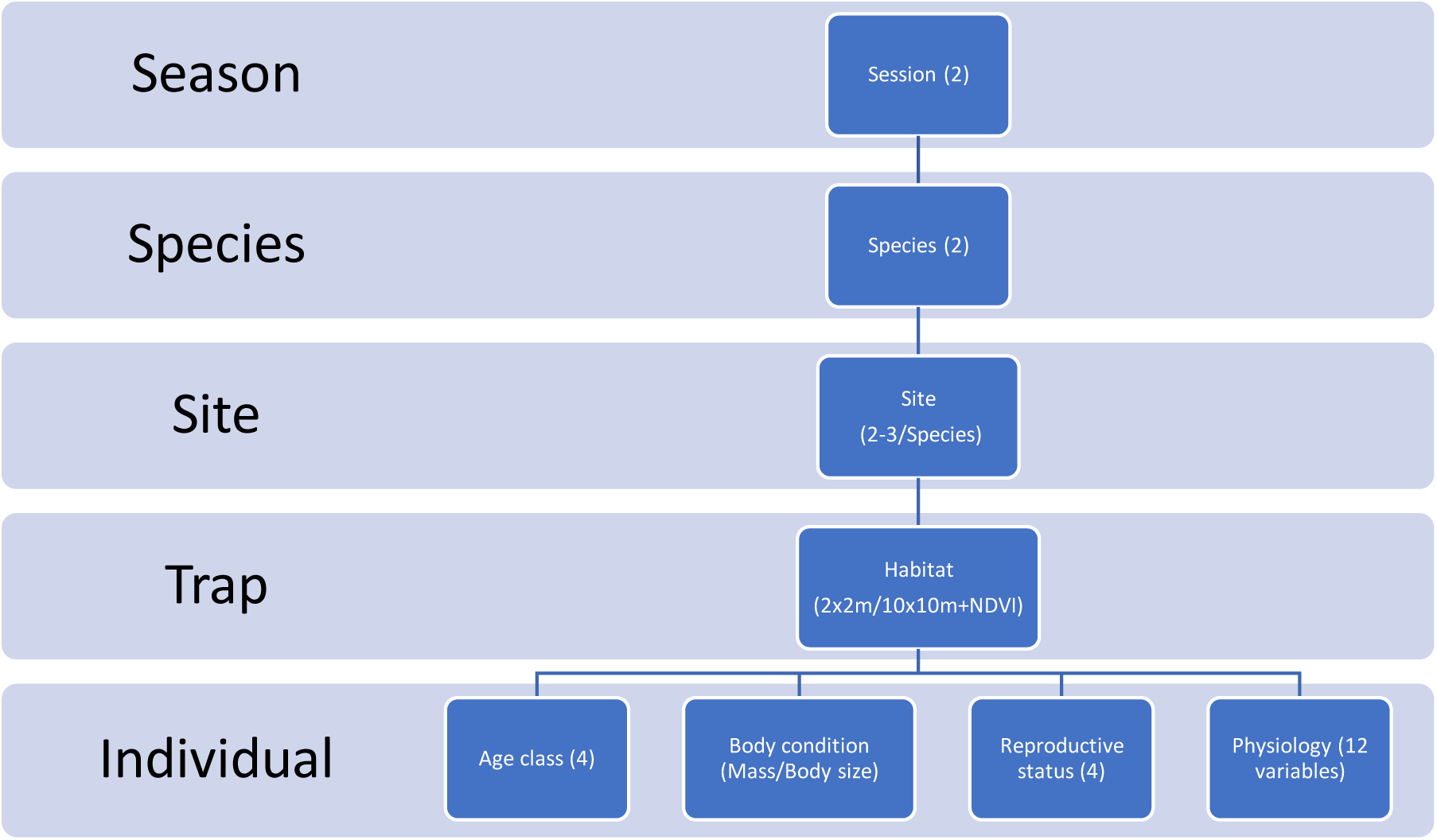
Graphical summary of organization and composition of data used for analysis in this study. Figures between brackets indicate number of modalities or types of categories included in the factor.

**Figure 3:**
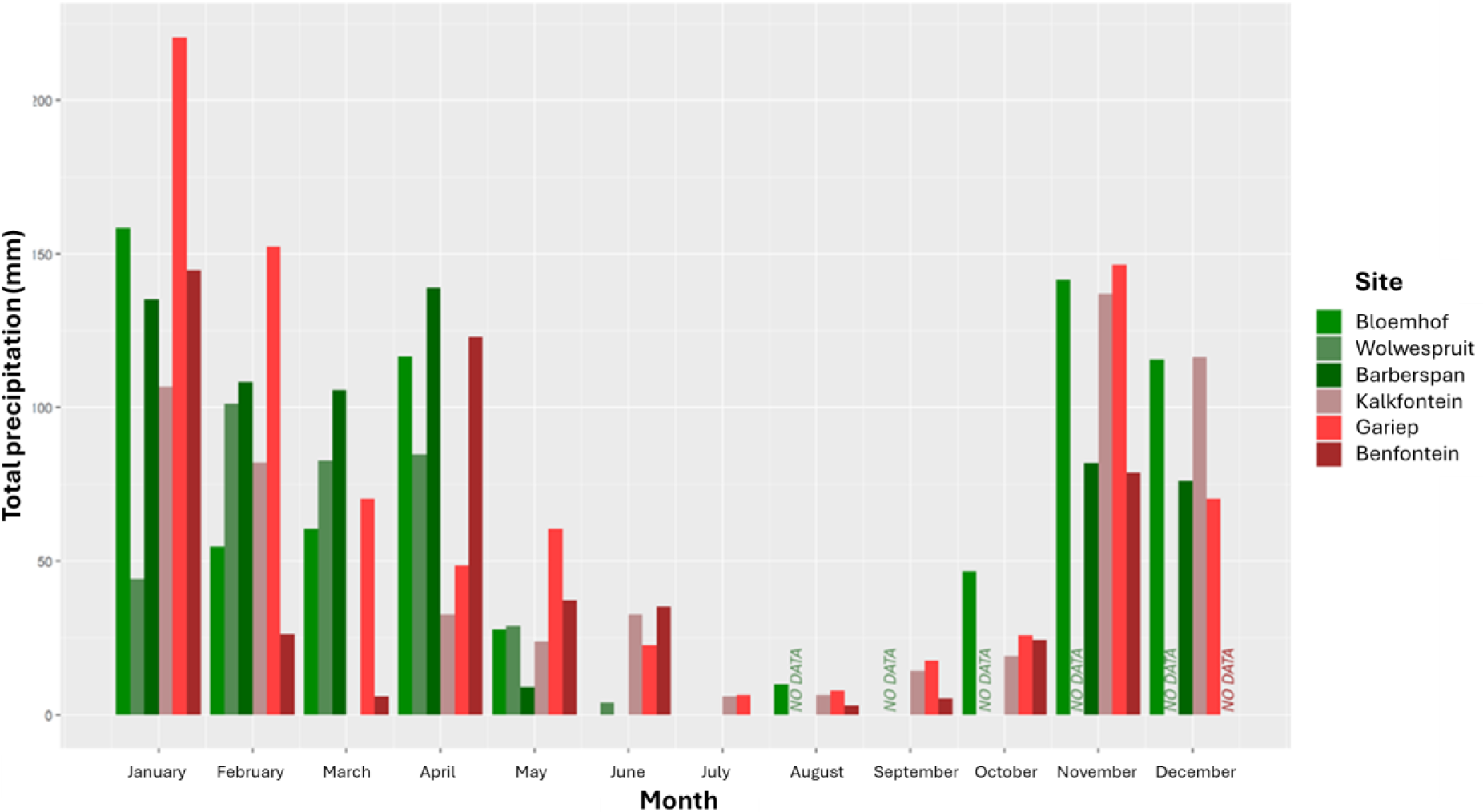
Total monthly precipitation for all sampled sites as recorded in 2022.

**Figure 4:**
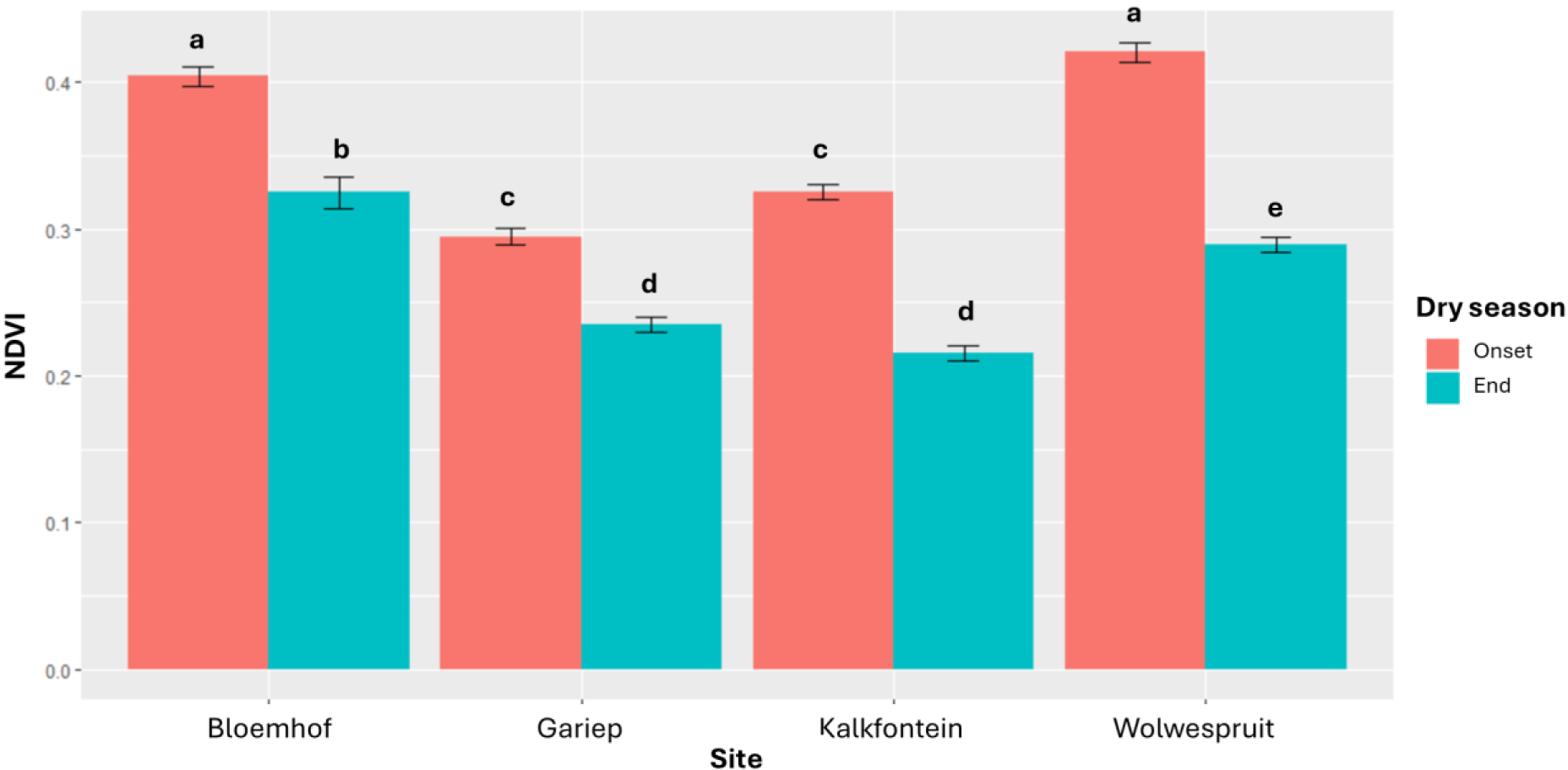
Average Normalized Differential Vegetation Index calculated within a buffer circle of 60m radius around each successful trap per site and session. NDVI was retrieved from the Copernicus Open Access Data Hub (Copernicus Sentinel-2 data [2023]). For each site and session, we retained NDVI data available for the closest day to the beginning of a sampling session, expected to represent the conditions experienced by the mice at the time of capture.

#### NDVI

The log-transformed NDVI of sites sampled at the two sessions (2 sites per species) was also compared between sites and sessions using *ANOVA*. Site, session and the interaction between these two variables (all p<0.001) contributed significantly to explain the NDVI variation (**Table 3**). A post-hoc Tukey multiple comparisons of means test showed that all sites had a significantly lower NDVI in September compared to May (**Figure 4**), indicating a poorer state of the vegetation at the end of the dry season. Furthermore, while these study locations were chosen in a region with similar semi-arid bioclimatic features, Gariep Dam and Kalkfontein (where *R.bechuanae* was targeted) seemed to stand out as having a lower NDVI than all other sites sampled in both May and in September, with Wolwespruit and Bloemhof (where *R.d.dilectus* was targeted) having the highest NDVI (**Figure 4**). Therefore, to take into account the impact of variations in NDVI between the two species trapping locations we included site as a random factor in analyses addressing body and physiological conditions.

**Table 3:** Results of the Analysis of Variance (ANOVA) test addressing variation in NDVI.

| Variable | Df | R<sup>2</sup> | F | P |
| --- | --- | --- | --- | --- |
| Session | 1 | 0.445 | 442.757 | **< 2.2\*10<sup>-16</sup>** |
| Site | 3 | 0.348 | 115.385 | **< 2.2\*10<sup>-16</sup>** |
| Session*Site | 3 | 0.021 | 7.113 | **1.512\*10<sup>-4</sup>** |
| Residual | 229 | 0.185 |  |  |

### Variation of Body condition

A linear mixed model assessed the influence of session, species, breeding status, age class and NDVI on body condition, including site as a random factor. The most parsimonious model included sex (p<0.001), age class (p<0.001), breeding status nested within sex (p<0.01) and the interaction between breeding status and session (p<0.05); session was not significant by itself p=0.965, and neither was the interaction between session and sex (p=0.683), while the effect of NDVI verged on significance (p=0.067, **Table 4**). Body condition was lower at the end of the dry season than at its onset for non-breeding females compared to breeding females (post-hoc pairwise t-test p<0.05, **Supplementary Table 10**).

**Table 4:** Results of the Analysis of Variance (ANOVA) test addressing variation in body condition.

| Variable | Df | R<sup>2</sup> | F | P |
| --- | --- | --- | --- | --- |
| Session | 1 | 1.271*10<sup>-5</sup> | 0.0019 | 0.965 |
| Sex | 1 | 0.085 | 25.163 | **1.072\*10<sup>-6</sup>** |
| NDVI | 1 | 8.007*10<sup>-4</sup> | 0.239 | 0.626 |
| Age class | 3 | 0.091 | 8.990 | **1.201\*10<sup>-5</sup>** |
| Breeding status | 2 | 0.041 | 6.067 | **0.003** |
| Session*Sex | 1 | 5.592*10<sup>-4</sup> | 0.167 | 0.683 |
| Session*Breeding status | 2 | 0.0299 | 4.444 | **0.012** |
| Residual | 229 | 0.753 |  |  |

### Variation in physiological responses

A *PERMANOVA* was performed using the distance matrix of 12 physiological markers as response variables and session, species, sex, breeding status, body condition, age class, NDVI and site (random factor) as explanatory variables (**Supplementary table 11**).

The most parsimonious model **(Table 5)** indicated an overall difference in physiological parameter concentrations between *R.bechuanae* and *R.d.dilectus* (p=0.002), as well as an influence of body condition (p=0.019), breeding status (nested within sex) (p=0.028), that differed between the two species (interaction effect, p=0.022), and the interaction between sex and species (p=0.020). Session (p=0.065), and sex (p=0.084) verged on overall significance and were also included in the model, indicating that physiological markers overall tend to shift between the start and end of the dry season, and to vary between sexes. Importantly, both species showed the same pattern of variation between sessions (no significant species*session interaction effect, p=0.498, **Supplementary Table 11**).

**Table 5:**
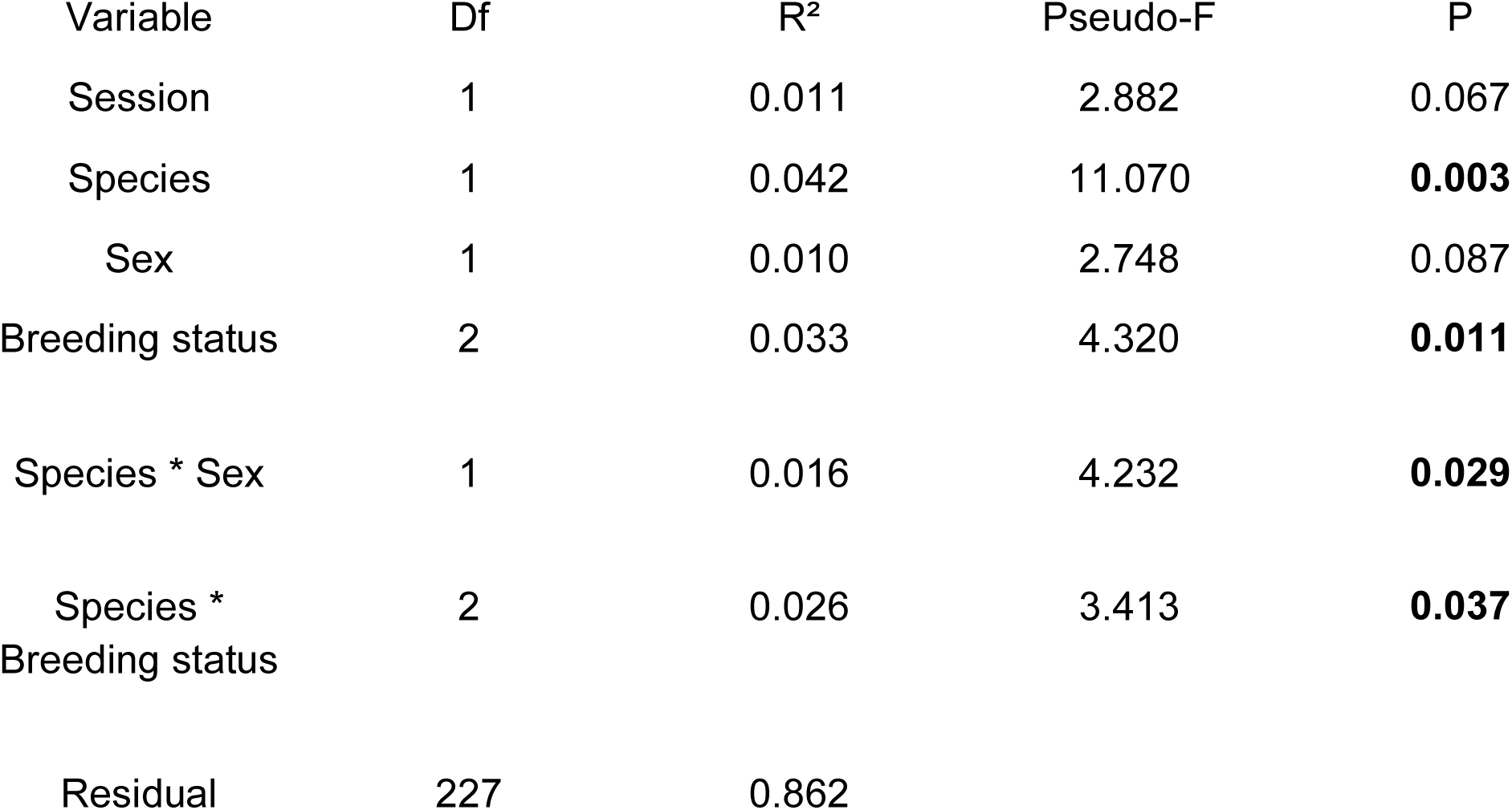
Summary of the results of the best fitting Permutational multivariate analysis of variance (PERMANOVA) model, addressing variation in physiological response (site= random factor)

Six physiological markers showed significant differences between the start and the end of the dry season (**Figure 5**). Blood concentrations of markers of nutrition and liver function (albumin, potassium, alanine aminotransferase, alkaline phosphatase, and total bilirubin) were lower at the end compared to the onset of the dry season (**Table 4**). Glucose also showed that trend, but not significantly (p=0.0577). In contrast, globulin levels varied in the opposite direction (higher at the end compared to the onset of the dry season).

**Figure 5:**
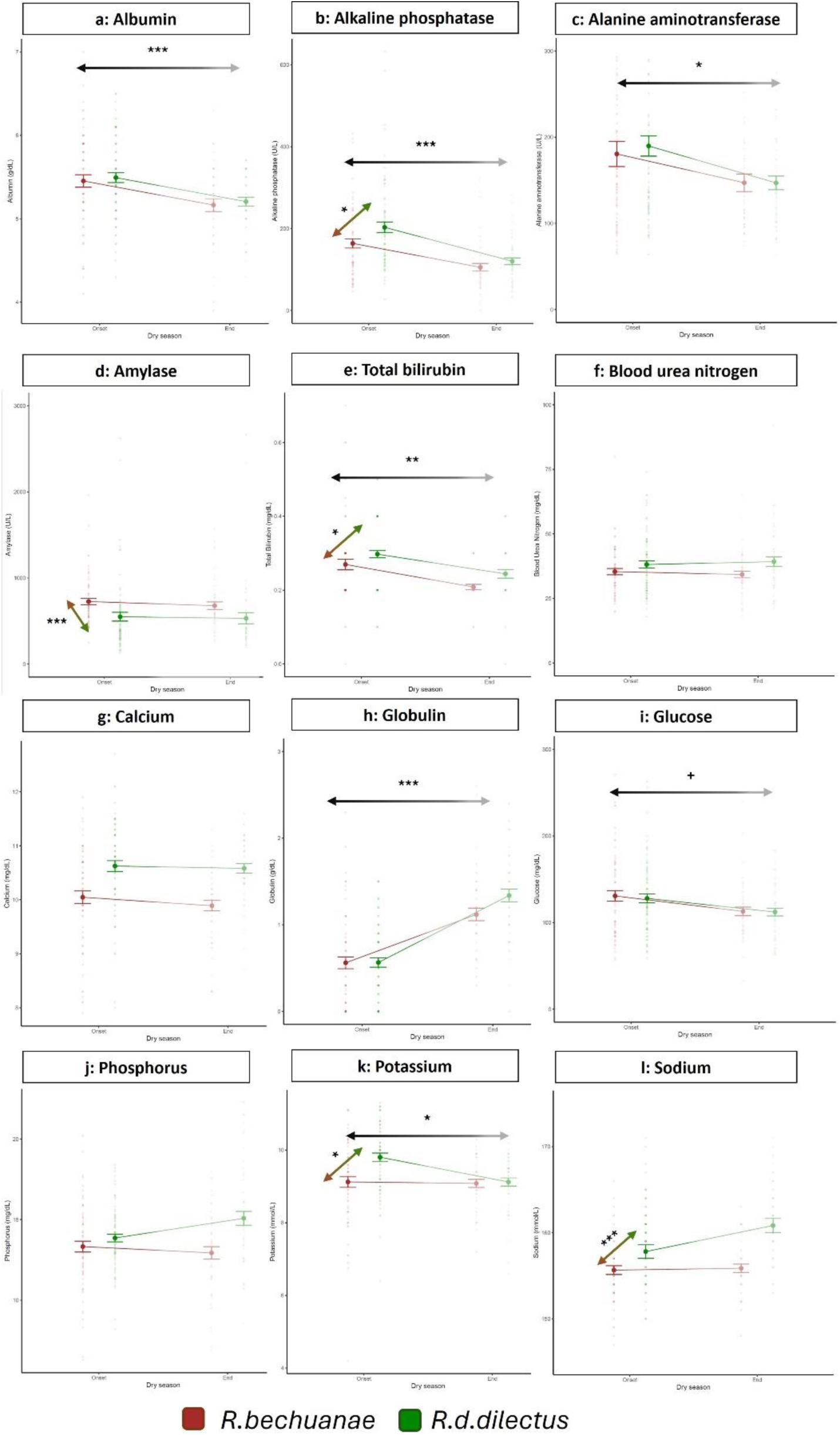
Levels of 12 physiological markers at the onset and end of the dry season in *R.bechuanae* and *R.d.dilectus*. Arrows indicate significant effects (p < 0.05) between the onset and the end of the dry season (black to grey gradient) or between the two species (brown to green gradient) following Bonferroni correction, with: **\*\*\***: p < 0.001; **\*\***: 0.001 < p < 0.01; **\***: 0.01 < p < 0.05; ^+^ : 0.05 < p < 0.1

The levels of four markers were significantly lower in *R.bechuanae* than in *R.d.dilectus* : total bilirubin, alkaline phosphatase (ALP), sodium and potassium (**Table 6**, **Figure 6**). However, amylase showed the opposite pattern between the species.

**Figure 6:** Significant interspecific differences involving four blood physiological marker concentrations (mean ± SE)

**Table 6:** Summary of the results of univariate mixed models addressing variation in responses of each of the 12 studied physiological variables (best fitting models).

| Physiological marker | Variable(s) kept in Best Model and their significance | adjusted R<sup>2</sup> | F | df Model | df Residual | P (model) |
| --- | --- | --- | --- | --- | --- | --- |
| Albumin | **Session<sup>\*\*\*</sup>, Sex<sup>\*</sup>** | 0.098 | 13.72 | 2 | 233 | 2.338*10<sup>-6</sup> |
| Alkaline phosphatase | **Session<sup>\*\*\*</sup>, Species<sup>\*</sup>, Sex<sup>\*\*\*</sup>, Breeding status<sup>+</sup>** | 0.260 | 17.43 | 5 | 229 | < 1.281*10<sup>-14</sup> |
| Alanine aminotransferase | **Session<sup>\*</sup>, Species, Sex, Breeding status, Species\*Sex, Species\*Breeding Status** | 0.060 | 2.884 | 8 | 227 | 0.004 |
| Amylase | **Species<sup>\*\*\*</sup>, Sex, Breeding status, Species\*Sex, Species\*Breeding Status** | 0.185 | 8.603 | 7 | 228 | 3.34*10<sup>-9</sup> |
| Calcium | Species | 0.009 | 3.095 | 1 | 234 | 0.080 |
| Globulin | **Session<sup>\*\*\*</sup>** | 0.291 | 97.44 | 1 | 234 | < 2.2*10<sup>-16</sup> |
| Glucose | Session<sup>+</sup>, **Sex<sup>\*\*</sup>** | 0.079 | 11.14 | 2 | 233 | 2.397 *10<sup>-5</sup> |
| Potassium | **Session<sup>\*</sup>, Species<sup>\*</sup>** | 0.068 | 9.569 | 2 | 233 | 1.014*10<sup>-4</sup> |
| Sodium | **Species<sup>\*\*\*</sup>, Sex** | 0.092 | 12.87 | 2 | 233 | 4.983*10<sup>-6</sup> |
| Blood Urea Nitrogen | *Null Random effect model* | 0.072 | / | / | / | / |
| Phosphorus | Species, Sex, **Breeding status<sup>\*</sup>** + random | 0.248 | / | / | / | ($\chi^2$ test vs. null model)<br>3.471*10<sup>-7</sup> |

*Note: The full models, before backwards elimination, included session, species, body condition, sex,* *breeding status, sex\*species and breeding status\*species as variables, with breeding status nested* *within sex and site as a partially-crossed random factor nested within species.*
**Bold:** significant effects ( $p < 0.05$ ) following Bonferroni correction with:
<sup>\*\*\*</sup>: $p < 0.001$ ; <sup>\*\*</sup>: $0.001 < p < 0.01$ ; <sup>\*</sup>: $0.01 < p < 0.05$ ; <sup>+</sup>: $0.05 < p < 0.1$

Regardless of species, the levels of glucose and ALP varied significantly with sex, tending to be lower in females than in males (**Supplementary figure 7**).

## Discussion

We asked how seasonal variation of dry conditions influenced the physiological response of semi-arid parapatric populations of two closely related *Rhabdomys* species that evolved under distinct environmental conditions (du Toit et al., 2012). Consistent with our predictions, we found strong evidence for physiological divergence between the species. We also observed a tendency for seasonal variation of the physiological response in both species related to energetic resource depletion at the end of the dry season, supported by the shifts in the levels of 6 blood parameters between the onset and the end of the dry season.

Our study took place in a relatively wet year (La Niña). Vegetation growth and senescence usually follow seasonal rainfall patterns; above-average rainfall widens the temporal window for vegetation growth, improving habitat conditions at the start of the typical dry season (Anyamba et al., 2002). Despite this, and consistent with our predictions, milder dry conditions did alter habitat quality of our study sites (as attested by shifts in NDVI and habitat structure).

### Seasonal variation of physiological condition

Despite our study year being relatively wet, mild dry conditions impacted blood marker concentrations in *R.bechuanae* and *R.d.dilectus*. These changes involved 6 out of 12 tested physiological markers, and included markers expected to vary in periods of dietary restriction, such as albumin. Contrary to our prediction, we did not find a significant interactive effect between session and species, suggesting that the semi-arid populations of the two species were similarly affected by seasonal dry conditions. However, a follow up study during an El Niño year may be necessary before one can conclude.

Water and nutritional stress are expected to impact the body condition of individuals (Fuller et al., 2021). As expected, body condition varied with breeding status, with differences between breeding and non-breeding females heightened at the end of the dry season. Additionally, several physiological markers varied significantly between males and females, indicating their association with physiological processes involved in reproduction. Reproduction in mammals has direct physiological costs, associated with increased energy, protein and calcium demands and indirectly through compensatory effects such as reduction in thermogenesis, immune function and physical activity (Speakman, 2008; Rintoul & Bringham, 2014; Stawski & Rojas, 2016; Schoepf et al., 2017b). It follows that sex-specific differences in reproductive investments are associated with contrasts in body weight, fat distribution and energy metabolism (Chen et al., 2012). In support, our results indicate lower levels of blood glucose, and a lower body condition in females compared with males. Moreover, albumin (ALB) and alkaline phosphatase (ALP) were significantly elevated in males compared to females, which could also be attributable to the pleiotropic effects of sex hormones (Havill et al., 2004; Sullivan et al., 2007).

The two species responded similarly to increased dryness while maintaining body condition, a proxy of fat reserves. Similarly, *R.pumilio* individuals tended to maintain their body condition stable while reducing their energy expenditure and physical activity during periods of limited food availability (Rimbach, Blanc et al., 2018; Rimbach, Jäger et al., 2018). The semi-arid populations of *R.bechuanae* and *R.d.dilectus* could be displaying a similar behavioral strategy.

Blood protein levels are generally good indicators of physiological condition (Tothova et al., 2016). They comprise two distinct and major components: albumin (ALB) and globulins (GLOB). In our study, ALB was significantly lower at the end than at the start of the dry season. ALB is synthesized in the liver and involved in the transport of bilirubin and several hormones. Blood ALB levels are less prevalent in malnourished individuals (Walker et al., 1990), supporting the hypothesis of reduction of quality and/or availability of food at the end of the dry season. The Nubian ibex *Capra nubiana*, a desert mammal, also displays low ALB during the dry season (Al-Eissa et al., 2012), which could reflect poor nutritional conditions, kidney dysfunction leading to poor water balance and high osmolality, or liver dysfunction causing low ALB synthesis. We found that GLOB was significantly higher at the end than at the start of the dry season. Higher GLOB levels were also reported for *R.pumilio* individuals that did not survive the dry season (Schoepf et al., 2017). GLOB is a family of molecules including immunity proteins and blood protein carriers (including transferrin). An increase in GLOB levels usually indicates elevation of immunoglobulins and hence could suggest heightened immune activity. Greater antibody production occurs in response to a higher prevalence of respiratory infections during the dry season, e.g. in the Nubian goat (Abdelatif et al., 2009), and/or reduced protein nutritional status in young birds (Lochmiller et al., 1993). An increase in blood GLOB concentrations may also reflect high levels of transferrin (Walker et al., 1990), a major GLOB that may accumulate in the blood in cases of severe iron deficiency (Kasvosve & Delanghe, 2002). Hence, elevated blood GLOB levels at the end of the dry season may indicate either or both depletion in iron resource (transferrin) and higher vulnerability to infection (immunoglobulin), both resulting from or being aggravated by reduced access to food. Lower levels of alkaline phosphatase (ALP), a liver enzyme, at the end of the dry season in *Rhabdomys*, is also evidence of malnutrition and deficiency in essential nutrients (proteins, magnesium, zinc) (Yousef et al., 2002; Saraç & Saygili, 2007; Ray et al., 2017). Alanine aminotransferase (ALT), another marker of liver function, also occurred at significantly lower concentrations at the end compared to the start of the dry season, indicating a reduction in liver activity (Walker et al., 1990) or malnutrition (Le Couteur et al., 2010). Indeed, as environmental resources become scarce, essential nutrients, such as pyridoxine (or vitamin B6), that constitute some ALT coenzymes, may be less available, leading to a reduction of ALT blood levels (Vespasiani-Gentilucci et al., 2018).

Blood glucose, another major physiological marker of nutrition state, was also slightly diminished at the end of the dry season, albeit not significantly. Fasting or low-energy feeding results in lower blood glucose levels (Jensen et al., 2013).

Acute starvation also typically causes bilirubin (TBIL) to accumulate in the blood, as the enzymatic process for conversion of heme into TBIL is stimulated (Thaler et al., 1972). Yet, TBIL was lower at the end compared to the onset of the dry season, rejecting the starvation hypothesis. Lower TBIL concentrations at the end of the dry season in *Rhabdomys* could stem from low hemoglobin counts because the body produces fewer red blood cells than usual; low hemoglobin can be induced by several factors, among which dietary factors such as iron deficiencies (Clark, 2008). Iron deficient anemia is also supported by higher GLOB concentrations observed at the end of the dry season. Infection can likewise be a cause of both decreased TBIL levels (Zhao et al., 2019) and higher GLOB concentrations at the end of the dry season. Finally, malnutrition and/or lack of potassium intake are among the main potential causes of the reduction of its levels in the blood (Their, 1986), which could explain why they were the lowest at the end of the dry season, when the available resources were lowest, in our study.

Overall, our results indicate that seasonal variation of some physiological markers could reflect reduced primary productivity throughout the dry season, which is coherent with our observation of habitat degradation (lower NDVI) at the end of the dry season. Blood concentrations of various markers seem to indicate difficulties in maintaining nutritional functions and the necessity of a higher immune output, caused or exacerbated by food restriction. Indeed, individuals undergoing malnutrition during dry conditions are expected to show low blood glucose concentrations, and deficiencies in essential nutrients, leading to pathologies such as anemia (Gordon et al., 1988). Indeed, animals foraging on soils potentially poor in nutrients such as iron, zinc, and magnesium are more prone to malnutrition (Graham, 1991; Gupta et al., 2008). Higher GLOB, lower TBIL and lower ALP blood concentrations in our study might relate to such deficiencies.

We did not observe seasonal variation of sodium or blood urea nitrogen, blood markers of kidney function or osmoregulation. Indeed, reduced environmental moisture at the end of the dry season could have caused an overall increase in blood osmolality, increasing blood concentrations of most biomarkers. Still, most markers exhibited a significant reduction in blood concentrations. This suggests that, compared to food availability, seasonal patterns in water availability may impose a milder physiological cost to the striped mice throughout the dry season in this semi-arid environment during a La Niña (wet) episode.

### Interspecific differences

*R.bechuanae* is thought to have evolved in drier environments than *R.d.dilectus*. Moreover, in the semi-arid region where they co-occur, we showed that *R.bechuanae* populations occupy slightly drier habitats than *R.d.dilectus*. While body condition did not differ significantly between species, some interspecific contrasts support our hypothesis that *R.bechuanae* may have developed a better physiological capacity to cope with dry conditions than *R.d.dilectus*.

Among the markers exhibiting a significant difference between *R.bechuanae* and *R.d.dilectus*, all but amylase showed higher blood concentrations in *R.d.dilectus*. Amylase is positively correlated with digestive activity and reflects differences in feeding status or diet composition between the species (Hidalgo et al., 1999). Given the role of alpha-amylase in digestion, lower blood amylase levels in *R.d.dilectus* than *R.bechuanae* may indicate lower digestive activity due to reduced access to food, or lower starch content in the diet. Pajic et al. (2019) found a correlation between dietary starch content and the number of amylase gene copies in several mammalian genomes, even in different habitats and with different diets. Since amylase activity and amylase gene expression are directly correlated with amylase gene copy number (Arendt et al., 2014), further studies could elucidate the potential genetic basis of the different levels of amylase in our two study species.

Sodium (NA) and potassium (K) concentrations, which depend on both dietary and water intake, were higher in *R.d.dilectus* than in *R.bechuanae*. These differences, coupled with higher ALP and TBIL concentrations in *R.d.dilectus* compared to *R.bechuanae*, could suggest a better nutrition state for *R.d.dilectus* than *R.bechuanae*. However, higher levels of amylase in *R.bechuanae* cast doubt on this interpretation. Instead, lower levels of blood NA, K, ALP, and TBIL in *R.bechuanae* would be more parsimoniously attributed to lower overall osmolality, perhaps due to better water balance in this species. A better overall ability of *R.bechuanae* to conserve body moisture could rely on physiological mechanisms such as more efficient excretion or osmoregulation, or through feeding on a diet optimizing water intake. We hypothesize that during the evolutionary history of *R.bechuanae*, selection pressures may have favored specific strategies allowing an overall more efficient water regulation compared to *R.d.dilectus*. These results, combined with the reduction in the levels of K, ALP and TBIL at the end of the dry season suggest both species are exposed to both water and nutrition stress.

We found relatively little evidence that the two species suffer differentially from lack of food in their semi-arid environment during a dry season impacted by La Niña phenomenon. Overall, energy levels, as indicated by blood glucose, and body condition, did not vary between the species. Instead, it seems that the overall differences between the two species can be accounted for by lower blood osmolality in *R.bechuanae*, and on average higher blood osmolality in *R.d.dilectus,* as evidenced by differences in NA and K concentrations, possibly due to overall poorer water regulation. High average blood amylase levels in *R.bechuanae*, compared to *R.d.dilectus* during the dry season, might indicate an interspecific difference in diet composition or food intake, rather than protein-energy malnutrition. Little is known about *R.bechuanae* and *R.d.dilectus* diets; the genus was characterised as a generalist feeding on seeds and other plant material, and insects (Curtis & Perrin, 1979). Some studies showed variation in diet composition within *Rhabdomys* genus associated with variation in local resources (Taylor & Green, 1976). The two species could have adjusted their diet to the respective local availabilities of different food resources within their home ranges. For example, during the dry season and in the semi-arid zone, arthropod biomass and diversity is affected by seasonal patterns in rainfall. Dalerum et al. (2017) found lower arthropod trapping success and diversity during the dry season, in shrub habitats compared to any other biomes, in Benfontein Game Reserve, one of our study sites. Thus, compared to seeds, the reliability of arthropods as a food source during dry conditions could be lower in shrublands, inhabited by *R.bechuanae* across its range and by both species in the semi-arid parapatry, than in grasslands inhabited by *R.d.dilectus* in allopatry. Variation in seed composition could also explain this interspecific difference in amylase concentrations. Indeed, granivores prefer seeds that have higher protein content and lower secondary metabolites but may be forced to take a wider range of seeds in dry areas than in mesic ones (Wilmer et al., 2009). If the hypothesis of a relatively ancient adaptation of *R.bechuanae* to dry habitats is true, it could be expected that it would also have adjusted its diet to the specific conditions encountered in dry habitats.

Within this ecosystem, *R.bechuanae* could forage more efficiently compared with *R.d.dilectus*, which evolved in a more mesic environment; its diet could also be richer in starch (such as containing more seeds/grains than leaves or fibre). Higher starch intake could help maintaining water balance, starch being the best substrate for metabolic water production in a dry environment (Adolph, 1964). This is consistent with the other interspecific differences revealed in this study, as diet can also significantly affect osmoregulation (Sabat et al., 2009).

## Perspectives and conclusions

Due to climate change, populations of many species experience changes in their habitats affecting their fitness and inducing shifts in their geographical distribution (Parmesan et al., 2006; Fuller et al., 2010). In some species, range shifts are expected to swell edge populations, as marginal habitats grow more favourable. For instance, edge populations of *Wilsonia citrina* hooded warblers have repeatedly expanded their ranges due to climate change (Melles et al., 2011). In other species, the quality of marginal habitats occupied by populations at the range edge may decline because of reduced food availability or nutrient richness, making these populations particularly vulnerable to climate change and inducing range contraction. For example, the koala *Phascolarctos cinereus* experiences declines in population sizes and habitat contraction due to droughts (Seabrook et al., 2011). As such, depending on their phenotypic plasticity, edge populations may hold the key to the survival of a species in a changing environment (Wu & Seebacher, 2022, Usui et al., 2023). *R.d.dilectus* edge populations may experience the same situation, as the example above, in the semi-arid zone, an hypothesis that could be tested addressing these populations dynamics, compared to *R.bechuanae* semi-arid populations.

In this study, we found that two closely related species with different ecological characteristics displayed similar seasonal negative shifts in blood concentrations of markers of nutrition and liver function in spite of a relatively mild seasonal drying of their habitats. We hence expect the physiological impact of harsher dry conditions, to affect the individuals’ fitness even more dramatically. The fact that the habitat of *R.bechuanae* studied populations was the driest at both time points, combined with our findings concerning their overall better water balance and nutrition than *R.d.dilectus*, point towards *R.bechuanae* coping better in the semi-arid environment during the dry season. Though adaptation *per se* cannot be inferred, this comparative study considering populations of the two species in the same biogeographic zone adds to the growing evidence that these cryptic species of striped mice have diverged phenotypically. Overall, these findings show the ecological relevance of physiological approaches in investigating the effect of climatic variation on organisms and question resilience of core populations of arid species and edge populations of mesic species in the face of predicted enhanced aridification.

## Acknowledgements

We thank R.Nokha who provided valuable and substantial field work contribution to this project. Constructive and insightful comments were provided by C.Smadja and C.Schradin helped improve a previous version of this paper. Technical support by D.Greuet (INM), L.Paradis (Geomatics, ISEM) and GenSeq platform. We are grateful to the rangers and personnel of Barberspan Bird Sanctuary, Benfontein Nature Reserve, Bloemhof Dam Nature Reserve, Gariep Dam Nature Reserve, Kalkfontein Dam Nature Reserve and Wolwespruit Nature Reserve who facilitated our work in these locations. Funding: CNRS/NRF & OSU-OREME

## Author contributions

G.G. conceived the idea and designed methodology with input from H.K; G.G., H.K., P.C., collected the data; H.K. analyzed the data with input from G.G.; H.K. wrote the first draft of the manuscript with input from G.G.; N.P. and N.A. contributed critically to the manuscript, secured funding for the project with G.G. and helped with field logistics. All authors approved the final version of the manuscript.

## Conflict of Interest statement

The authors declare they have no conflict of interest.

## Notes

### Competing Interest Statement

The authors have declared no competing interest.

### Summary of Updates

Link to PCI peer review recommendation, as well as link to the data and scripts used in this study

