## Supplementary Material for "Negative impact of mild arid conditions in natural rodent populations revealed using markers of physiological condition in natura"

**Supplementary Table 1 : trapping effort per site per day**

| **Site** | **Date** | **#traps** |
| --- | --- | --- |
| Gariep Dam | 04-mai | 192 |
|  | 05-mai | 432 |
|  | 06-mai | 408 |
|  | 07-mai | 240 |
|  | 08-mai | 232 |
| Kalkfontein | 09-mai | 59 |
|  | 10-mai | 389 |
|  | 11-mai | 360 |
|  | 12-mai | 157 |
|  | 13-mai | 4 |
| Benfontein | 14-mai | 293 |
|  | 15-mai | 192 |
|  | 16-mai | 74 |
| Bloemhof | 18-mai | 286 |
|  | 19-mai | 374 |
|  | 20-mai | 146 |
| Barberspan | 22-mai | 120 |
|  | 23-mai | 390 |
|  | 24-mai | 322 |
|  | 25-mai | 276 |
|  | 26-mai | 205 |
| Wolwespruit | 27-mai | 191 |
|  | 28-mai | 183 |
|  | 29-mai | 87 |
| Kalkfontein | 11-sept | 303 |
|  | 12-sept | 198 |
|  | 13-sept | 243 |
|  | 14-sept | 97 |
| Gariep Dam | 16-sept | 336 |
|  | 17-sept | 418 |
|  | 18-sept | 317 |
|  | 19-sept | 276 |
|  | 20-sept | 155 |
| Bloemhof | 22-sept | 331 |
|  | 23-sept | 394 |
|  | 24-sept | 337 |
|  | 25-sept | 282 |
|  | 26-sept | 122 |
| Wolwespruit | 27-sept | 127 |
|  | 28-sept | 209 |
|  | 29-sept | 76 |
|  | 30-sept | 133 |

**Supplementary Table 2 : Trapped surface per site and trapping session**

| **Site** | **Session** | **Trapped surface (km²)** |
| --- | --- | --- |
| Barberspan | May | 0,2185 |
| Benfontein | May | 0,1733 |
| Bloemhof | May | 0,1909 |
| Bloemhof | September | 0,1764 |
| Gariep | May | 0,0629 |
| Gariep | September | 0,0912 |
| Kalkfontein | May | 0,0493 |
| Kalkfontein | September | 0,0907 |
| Wolwespruit | May | 0,1646 |
| Wolwespruit | September | 0,1679 |

| **Sample** | **Rotor** | **ALB** | **ALP** | **ALT** | **AMY** | **TBIL** | **BUN** | **CA** | **PHOS** | **CRE** | **GLU** | **NA** | **K** | **TP** | **GLOB** |
| --- | --- | --- | --- | --- | --- | --- | --- | --- | --- | --- | --- | --- | --- | --- | --- |
| 13 | 1 | 4,8 | 114 | 122 | 1108 | 0,6 | 36 | 9,8 | 12,8 | 0,0 | 86 | 160 | 7,0 | 6,0 | 1,0 |
| 13 | 2 | 5,7 | 111 | 121 | 1026 | 0,3 | 32 | 10,6 | 12,6 | 0,0 | 83 | 164 | 6,8 | 6,4 | 0,7 |
| **%CV1** | | **12,12** | **1,89** | **0,58** | **5,43** | **47,14** | **8,32** | **5,55** | **1,11** | **0,00** | **2,51** | **1,75** | **2,05** | **4,56** | **24,96** |
| 15 | 1 | 6,2 | 253 | 88 | 699 | 0,2 | 38 | 11 | 14,6 | 0,3 | 115 | 158 | 8,9 | 6,1 | 0,0 |
| 15 | 2 | 6,4 | 236 | 90 | 714 | 0,2 | 39 | 10,9 | 15 | 0,3 | 113 | 158 | 9 | 6,2 | 0,0 |
| **%CV2** | | **2,24** | **4,92** | **1,59** | **1,50** | **0,00** | **1,84** | **0,65** | **1,91** | **0,00** | **1,24** | **0,00** | **0,79** | **1,15** | **0,00** |
| 18 | 1 | 5,6 | 158 | 127 | 878 | 0,2 | 33 | 10,2 | 11,7 | 0,0 | 169 | 155 | 8,4 | 5,8 | 0,2 |
| 18 | 2 | 5,8 | 174 | 121 | 912 | 0,3 | 33 | 10,2 | 11,8 | 0,0 | 185 | 156 | 8,5 | 6,1 | 0,3 |
| **%CV3** | | **2,48** | **6,82** | **3,42** | **2,69** | **28,28** | **0,00** | **0,00** | **0,60** | **0,00** | **6,39** | **0,45** | **0,84** | **3,57** | **28,28** |
| 35 | 1 | 6,1 | 152 | 66 | 420 | 0,2 | 24 | 11,4 | 12 | 0,0 | 237 | 155 | 9,9 | 6,7 | 0,6 |
| 35 | 2 | 6,1 | 162 | 68 | 427 | 0,2 | 25 | 11,5 | 12,2 | 0,0 | 233 | 158 | 9,9 | 6,7 | 0,6 |
| **%CV4** | | **0,00** | **4,50** | **2,11** | **1,17** | **0,00** | **2,89** | **0,62** | **1,17** | **0,00** | **1,20** | **1,36** | **0,00** | **0,00** | **0,00** |
| 52 | 1 | 4,3 | 85 | 75 | 815 | 0,4 | 43 | 9,9 | 6,4 | 0,3 | 101 | 154 | 6,6 | 5,9 | 1,5 |
| 52 | 2 | 4,5 | 89 | 75 | 832 | 0,2 | 45 | 10 | 6,6 | 0,0 | 101 | 158 | 6,9 | 5,9 | 1,4 |
| **%CV5** | | **3,21** | **3,25** | **0,00** | **1,46** | **47,14** | **3,21** | **0,71** | **2,18** | **134,84** | **0,00** | **1,81** | **3,14** | **0,00** | **4,88** |
| 103 | 1 | 4,7 | 81 | 541 | 165 | 0,4 | 25 | 8,1 | 11,1 | 0,0 | 121 | 127 | 10,1 | 5,9 | 0,0 |
| 103 | 2 | 4,7 | 95 | 524 | 164 | 0,4 | 25 | 8 | 11,3 | 0,0 | 122 | 129 | 8,9 | 5,9 | 0,0 |
| **%CV6** | | **0,00** | **11,25** | **2,26** | **0,43** | **0,00** | **0,00** | **0,88** | **1,26** | **0,00** | **0,58** | **1,10** | **8,93** | **0,00** | **0,00** |

**Supplementary Table 3 : Intra-sample coefficients of variation (CV) for 14 measured physiological markers**

ALB : Albumin (g/dL); ALP : Alkaline phosphatase (U/L); ALT : Alanine Aminotransferase (U/L); AMY : Amylase (U/L) ; TBIL : Total Bilirubin (mg/dL); BUN : Blood Urea Nitrogen (mg/dL); CA : Calcium (mg/dL); PHOS : Phosphorus (mg/dL); CRE : Creatinine (mg/dL); GLU : Glucose (mg/dL); NA : Sodium (mmol/L); K : Potassium (mmol/L); TP : Total Protein (g/dL); GLOB : Globulin (g/dL)

For these six samples, the calculated CV equaled 5.6 ± 1.9 %, which is consistent with the intra-test CV% provided in the manufacturer’s instruction manual for other mammal species (<10 %)

**Supplementary Methods**

**Aridity Index formula :**

The Aridity Index formula as used in this article was as follows: $AI=\frac{P}{PET}$

where PET is the potential evapotranspiration and P is the average annual precipitation (UNEP, 1992).

PET by Thornthwaite method (Thornthwaite, 1948) was calculated using monthly Temperature Ti, from which we first calculated a Heat index I_i_ as follows :

$$I_{i}={(\frac{T_{i}}{5})}^{1.514}$$

Temperature efficiency index *J* (sum of monthly heat indexes) was then calculated as follows:

$$J=\sum i=1^{12}(I_{i})$$

With exponent *c* :

$$c = 0.000000675J^{3} - 0.0000771J^{2}+ 0.01792J + 0.49239$$

Potential evapotranspiration at 0° Latitude was then calculated as follows :

$$\mathrm{PET}_{i} (0) = 1.6(10\frac{T_{i}}{J})c$$

Then, in order to take into account photoperiod, and Potential Evapotranspiration at the corresponding latitude L was obtained using K, a constant for each month of the year, varying with latitude:

$${PET}_{i} (L) = K{PET}_{i} (0)$$

This was the final value of average monthly potential evapotranspiration, added together to form annual potential evapotranspiration, over which annual precipitation was divided to obtain aridity index.

**Individuals excluded from the study after genotyping:**

As this study took place in a semi-arid region where the ranges of several species of *Rhabdomys* overlap, striped mice belonging to different species than the targeted ones were trapped in some reserves. However, they rarely shared habitats, such that trapping was mostly specific to the targeted species.

In total, nine individuals were excluded from the analysis because they belonged to *R.d.chakae*, a closely related subspecies to *R.d.dilectus* in Wolwespruit (n=2) and Barberspan (n=1), five *R.d.dilectus* individuals were trapped in Kalkfontein Dam Reserve and 1 in Gariep Dam Nature Reserve. The proportion of sampled individuals from the “correct” species before exclusion from the analyses in each site was respectively 92.31 % in Barberspan, 91.23 % in Kalkfontein, 100 % in Bloemhof, 100 % in Benfontein, 97.96 % in Gariep and 96.55 % in Wolwespruit.

**Scaled mass index formula :**

The scaled mass index of body condition (Peig & Green, 2009) was computed as follows :

**Scaled Mass Index** $SMI =M_{i}{[\frac{L_{0}}{L_{i}}]}^{b_{SMA}}$

Where $M_{i}$ and $L_{i}$ are the body mass and length of individual I, respectively; $b_{SMA}$is the scaling exponent estimated by the standardized major axis (SMA) regression or mass over length using R package *smatr* (v.3.4-8); $L_{0}$ is an arbitrary length value, here the mean length across all individuals.

**Permutational ANOVA and dispersion tests on habitat data:**

We compared the vegetation characteristics surrounding each successful trap with a *PERMANOVA*, using the first 5 Principal Component coordinates (**Supplementary Tables 7 & 9**) of each trap as response variable, and site and session as explanatory variables. For 4m² quadrats, PC1 was most explained by variance in dry grass (39.85 %), green bushes (35.63 %) and uncovered surfaces (20.59 %), PC2 was most explained by variance in the presence of green grass (62.04 %), PC3 by presence of holes in the ground (60.13 %), PC4 by presence of dry bushes (74.2 %) and PC5 by presence of succulent plants (76.56 %). For 100m² quadrats, PC1 was most explained by variance in dry grass (32.14 %), green bushes (26.16 %), uncovered surfaces (15.91 %) and dry bushes (10.96 %); PC2 was most explained by green grass (47.24 %), dry grass (20.85 %), holes (13.59 %), and uncovered surfaces (11.77 %); PC3 by succulent plants (53.34 %), PC4 by dry bushes (34.28 %), uncovered surfaces (28.32 %) and succulent plants (23.96 %); and PC5 by holes (43.92 %), green bushes (25.85 %) and succulent plants (18.69 %).

To assess differences in heterogeneity of habitat characteristics between sampling sites or

sessions, we also performed a permutational multivariate dispersion test (*permdisp*, package *vegan*, version 2.6-4) on the categorical explanatory variables. At the 2x2m level, the habitats were significantly more heterogeneous at the start compared to the end of the dry season (p=0.024), with additional inter-site differences (p<0.001, **Supplementary Table 5**). At the onset of the dry season, conditions were highly variable on PC2 (mostly associated with green grass characteristics), but much less so at the end of the dry season (**Supplementary Figure 1**), with more intermediate percentages values of green grass recorded in September. This could be explained by the fact that as the environment got drier, micro-habitats covered by large patches of green grass became less available, while our striped mice tended to avoid the driest areas, poor in both cover and resources. Among sites sampled in both sessions, habitat characteristics in Kalkfontein were significantly more heterogeneous than in Gariep and Bloemhof. Significant differences in dispersion were also found between sites at the 10x10m level, with conditions in Kalkfontein being again characterized by the highest dispersion (p=0.001, **Supplementary Table 6**); however, no significant difference was found between sessions.

**Permutational dispersion tests on physiological data:**

We performed a permutational multivariate dispersion test on all categorical explanatory variables. *PERMDISP* yielded no significant differences in dispersion patterns between groups except for “breeding status”. A post-hoc Tukey test indicated that the latter difference was due to higher dispersion in physiological characteristics of breeding (N=54) vs. non-breeding males (N=95). As PERMANOVA is generally robust to dispersion heterogeneity in slightly unbalanced designs (Anderson, 2013), we considered this as a true effect of the explanatory variable. In the subsequent univariate analyses, variance homoscedasticity was in each case carefully assessed in diagnostic plots of residuals *vs*. fitted values.

**Supplementary Results**

***Supplementary Table 4 :* Results of the permutational dispersion test (Habitat data, 2x2m quadrats)**

Explanatory variable: Session

| Variable | Df | F | P |
| --- | --- | --- | --- |
| Session | 1 | 5.139 | 0.024 |
| Residual | 234 |  |  |

Explanatory variable: Site

| Variable | Df | F | P |
| --- | --- | --- | --- |
| Session | 5 | 6.126 | 2.381*10^-5^ |
| Residual | 230 |  |  |

***Supplementary Table 5 :* Results of the permutational dispersion test (Habitat data, 10x10m quadrats)**

Explanatory variable: Session

| Variable | Df | F | P |
| --- | --- | --- | --- |
| Session | 1 | 1.317 | 0.252 |
| Residual | 234 |  |  |

Explanatory variable: Site

| Variable | Df | F | P |
| --- | --- | --- | --- |
| Session | 5 | 4.243 | 0.001 |
| Residual | 230 |  |  |

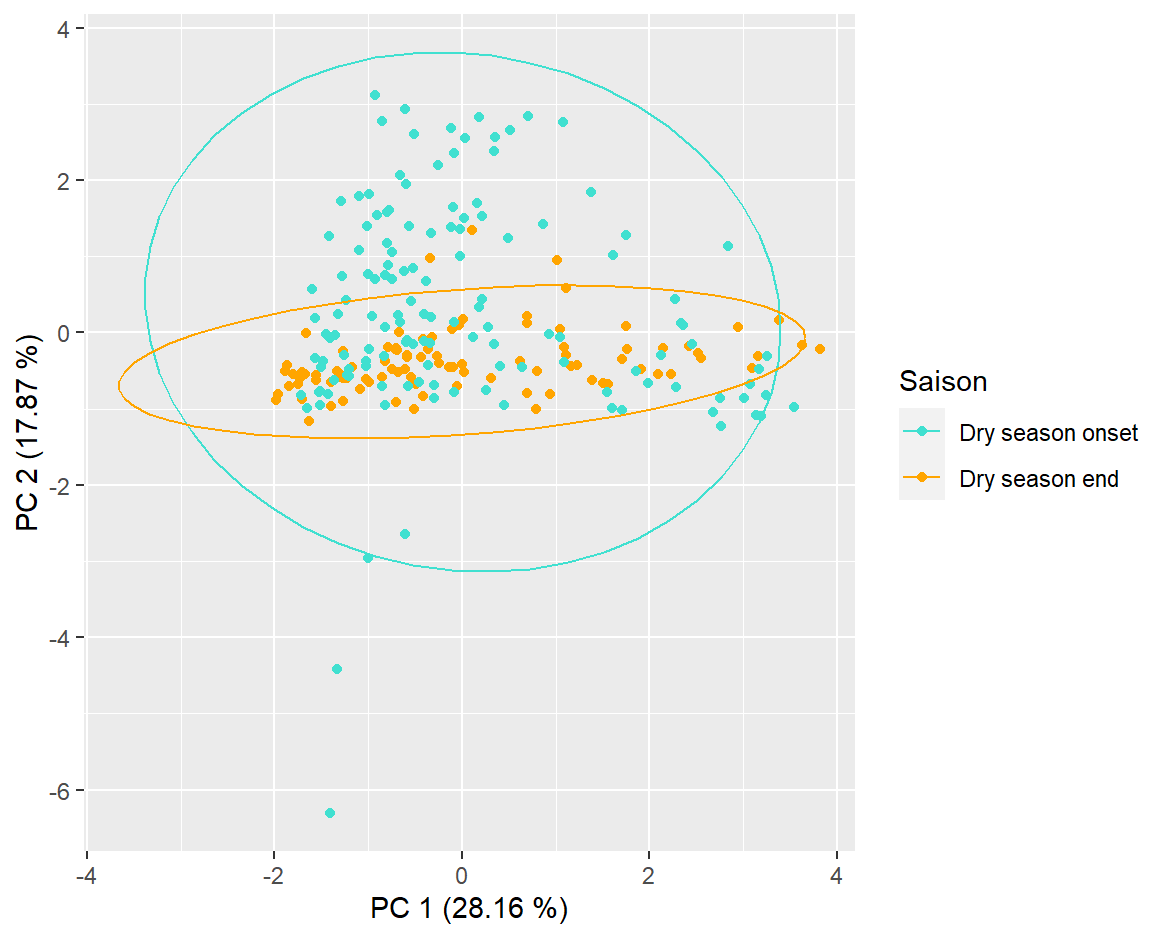

**Supplementary Figure 1: *First two axes of the principal Component Analysis plot identifying the position of traps in which 7 habitat structure variables were measured, for each sampling session (based on 2x2m quadrats)***

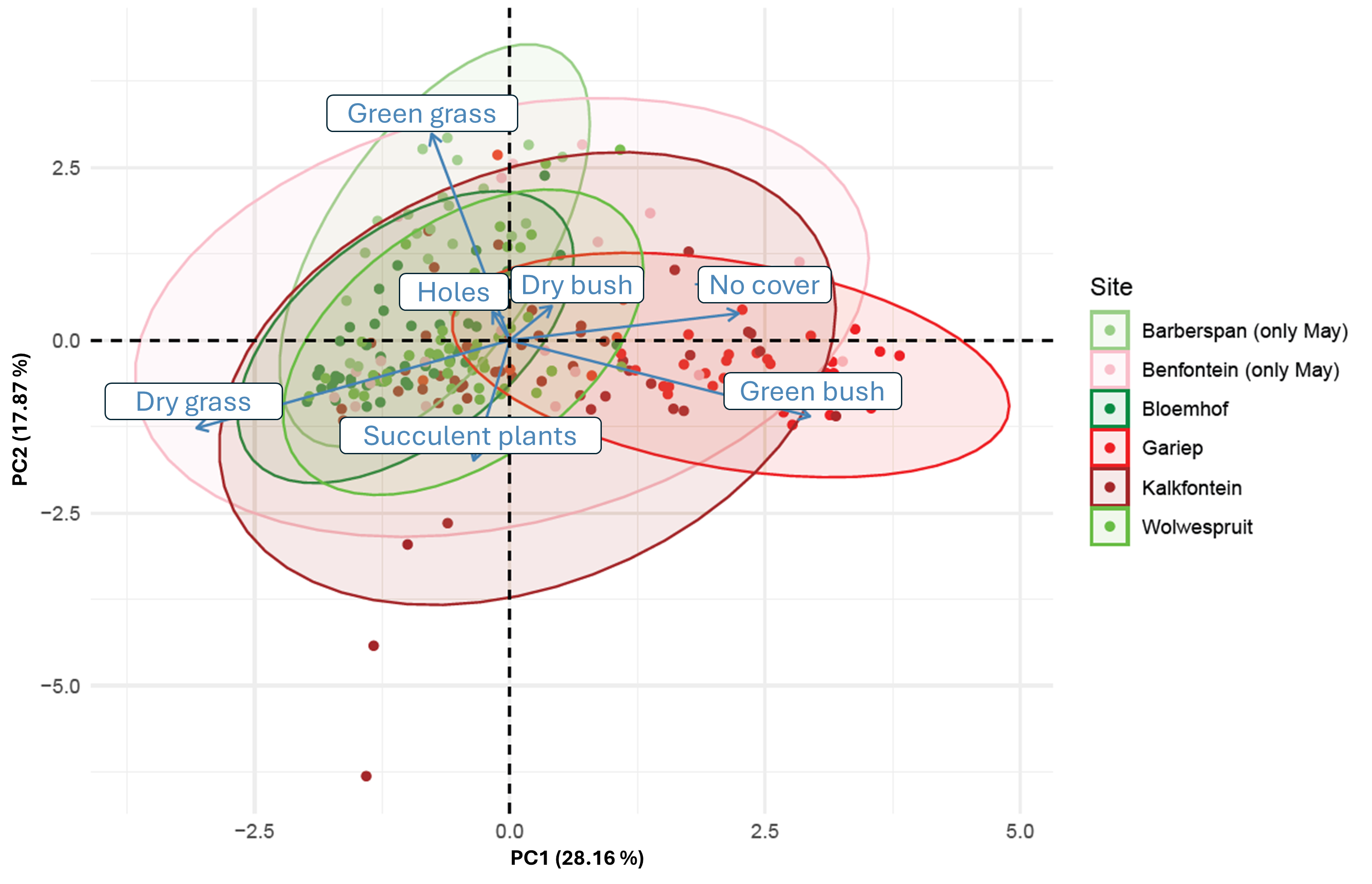

**Supplementary Figure 2: *First two axes of the principal Component Analysis biplot identifying the position of traps in which 7 habitat structure variables were measured, for each sampling site, as well as the position of 7 measured habitat structure variables (based on 2x2m quadrats)***

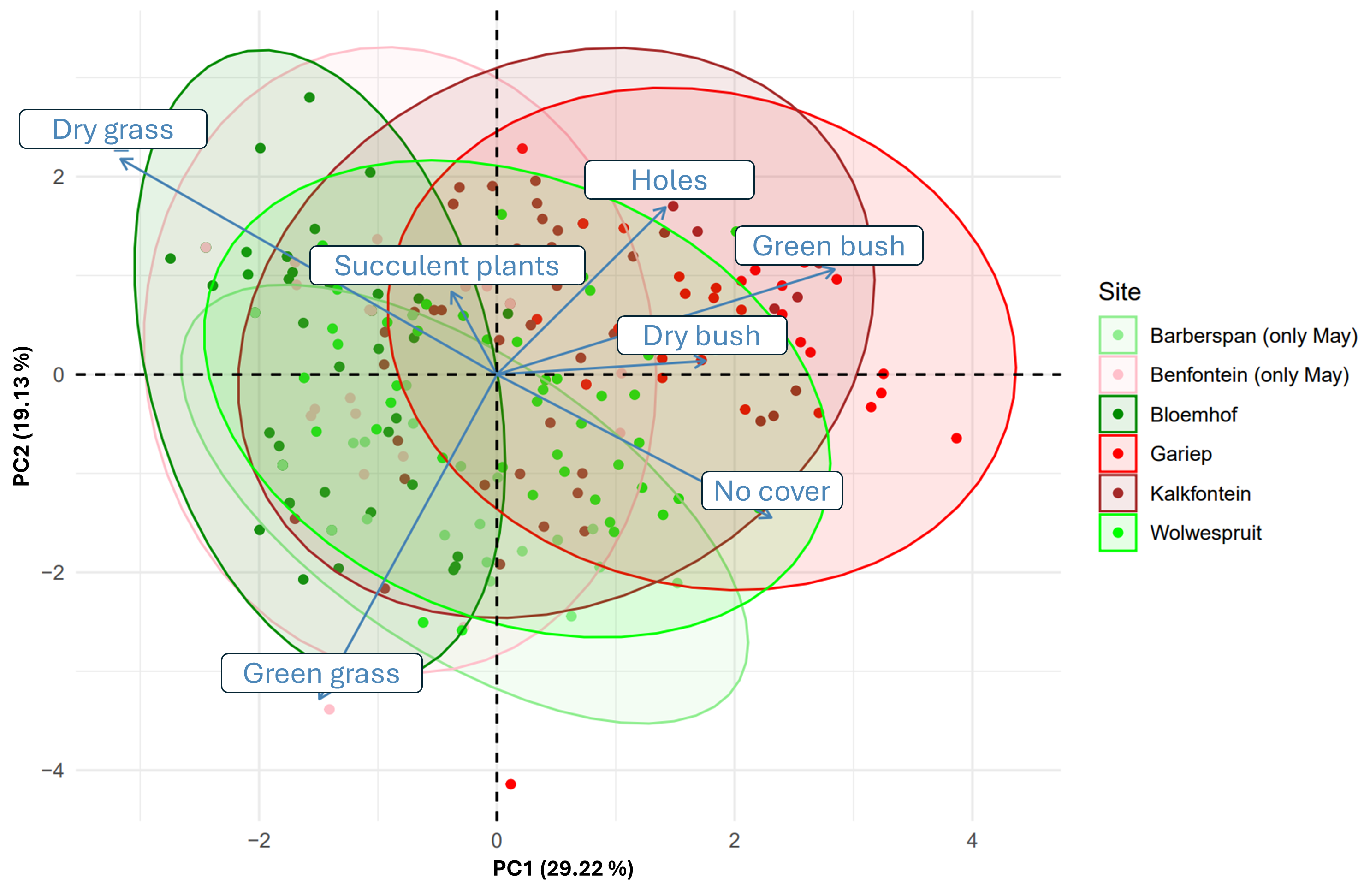

**Supplementary Figure 3: *First two axes of the principal Component Analysis biplot identifying the position of traps in which 7 habitat structure variables were measured, for each sampling site, as well as the position of 7 measured habitat structure variables (based on 10x10m quadrats)***

***Supplementary Table 6: Proportion of Variance explained by each principal component pictured in Supp.Fig.4***

|  | eigenvalue | percentage of variance | cumulative percentage of variance |
| --- | --- | --- | --- |
| PC 1 | 1.971 | 28.164 | 28.164 |
| PC 2 | 1.251 | 17.874 | 46.038 |
| PC 3 | 1.108 | 15.823 | 61.861 |
| PC 4 | 1.026 | 14.656 | 76.517 |
| PC 5 | 0.934 | 13.342 | 89.859 |
| PC 6 | 0.710 | 10.141 | 100.000 |
| PC 7 | 2.414 x 10^-30^ | 3.449 x 10^-29^ | 100.000 |

***Supplementary Table 7:* Results of the Permutational Multivariate Analysis of Variance (Habitat data, 2x2m quadrats)**

| Variable | Df | R² | Pseudo-F | P |
| --- | --- | --- | --- | --- |
| Session | 1 | 0.038 | 12.385 | 9.999*10^-5^ |
| Session/Site | 8 | 0.269 | 10.992 | 9.999*10^-5^ |
| Residual | 226 | 0.693 |  |  |

***Supplementary Table 8 : Proportion of Variance explained by each principal component pictured in Supp.Fig.5***

|  | eigenvalue | percentage of variance | cumulative percentage of variance |
| --- | --- | --- | --- |
| PC 1 | 2.045 | 29.216 | 29.216 |
| PC 2 | 1.339 | 19.132 | 48.348 |
| PC 3 | 1.036 | 14.804 | 63.152 |
| PC 4 | 0.944 | 13.488 | 76.640 |
| PC 5 | 0.933 | 13.323 | 89.963 |
| PC 6 | 0.702 | 10.030 | 99.993 |
| PC 7 | 4.680 x 10^-4^ | 0.007 | 100.000 |

***Supplementary Table 9: Permutational Multivariate Analysis of Variance table (10x10m vegetation quadrats)***

| Variable | Df | R² | Pseudo-F | P |
| --- | --- | --- | --- | --- |
| Session | 1 | 0.101 | 38.742 | 9.999*10^-5^ |
| Session*Site | 8 | 0.308 | 14.715 | 9.999*10^-5^ |
| Residual | 226 | 0.591 |  |  |

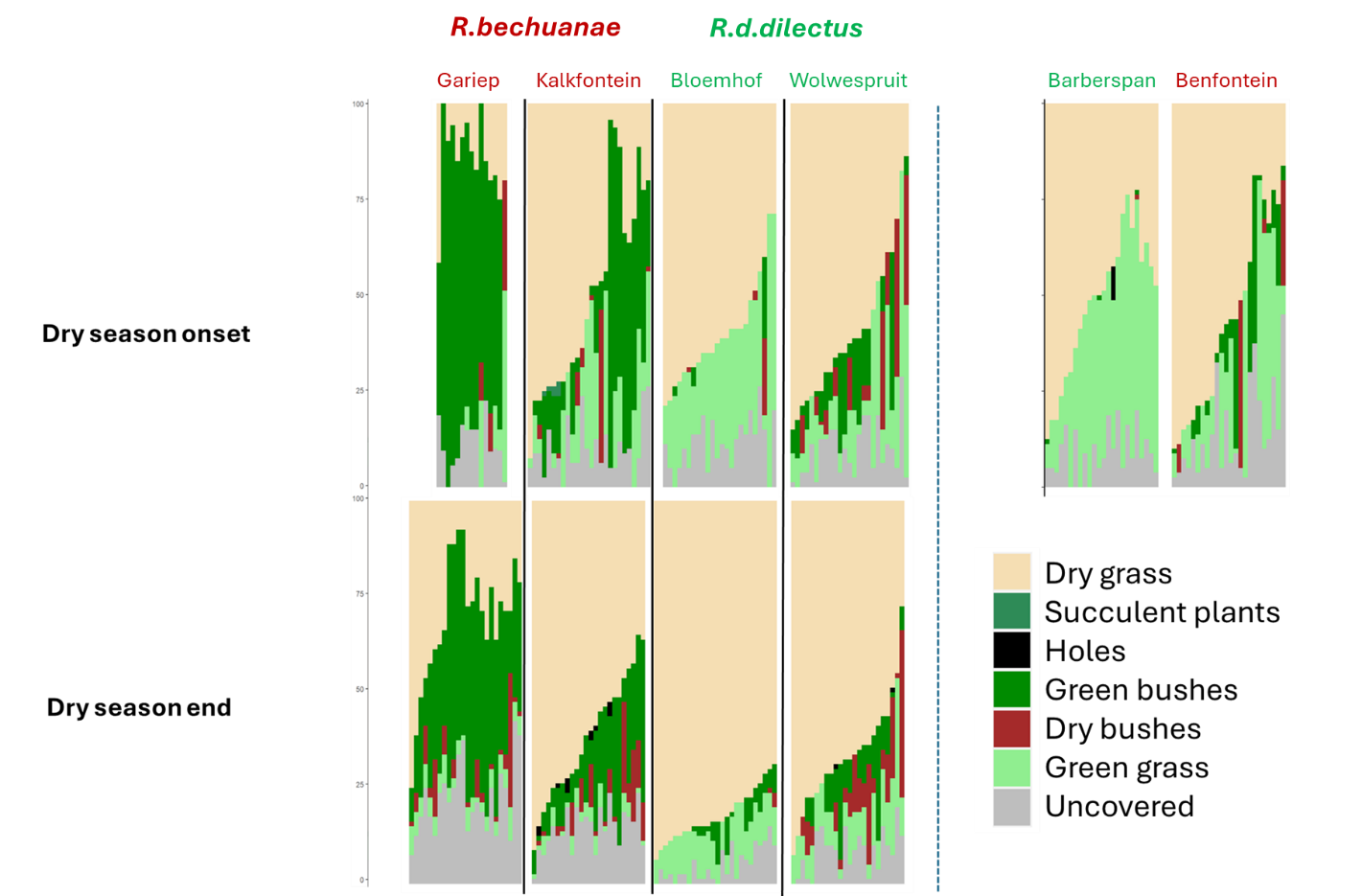

***Supplementary Figure 4 : Vegetation composition of each surveyed trap per site and sampling session (2x2m quadrats)***

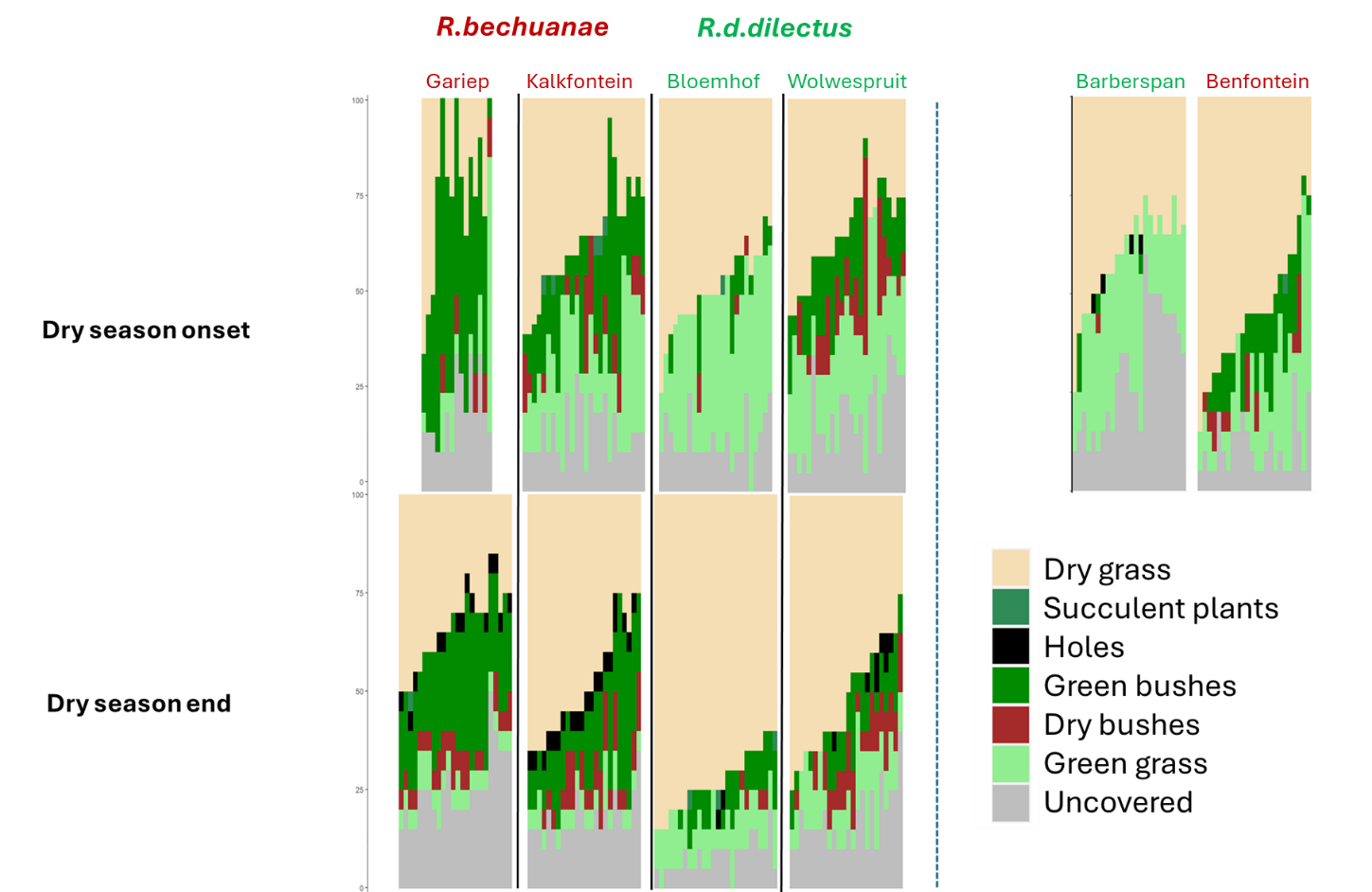

***Supplementary Figure 5 : Vegetation composition of each surveyed trap per site and sampling session (10x10m quadrats)***

**Supplementary table 10: Results of pairwise comparisons of body condition data using t-tests (p-value adjustment method: Bonferroni)**

Explanatory variable: Breeding status

|  | NBF | BF | NBM |
| --- | --- | --- | --- |
| BF | **< 0.001** | - | - |
| NBM | 0.0644 | - | - |
| BM | **-** | 0.4476 | 0.8084 |

NBF: Non-breeding female ; BF : Breeding female; NBM : Non-breeding male; BM : Breeding male

Interaction effect with Session

**Dry season onset**

|  | NBF | BF | NBM |
| --- | --- | --- | --- |
| BF | 1.000 | - | - |
| NBM | 1.000 | - | - |
| BM | - | 1.000 | 1.000 |

**Dry season end**

|  | NBF | BF | NBM |
| --- | --- | --- | --- |
| BF | **< 0.001** | - | - |
| NBM | 0.055 | - | - |
| BM | **-** | 1.000 | 1.000 |

**Inter-session contrasts**

| NBF | 0.146 |
| --- | --- |
| BF | 1.000 |
| NBM | 1.000 |
| BM | 1.000 |

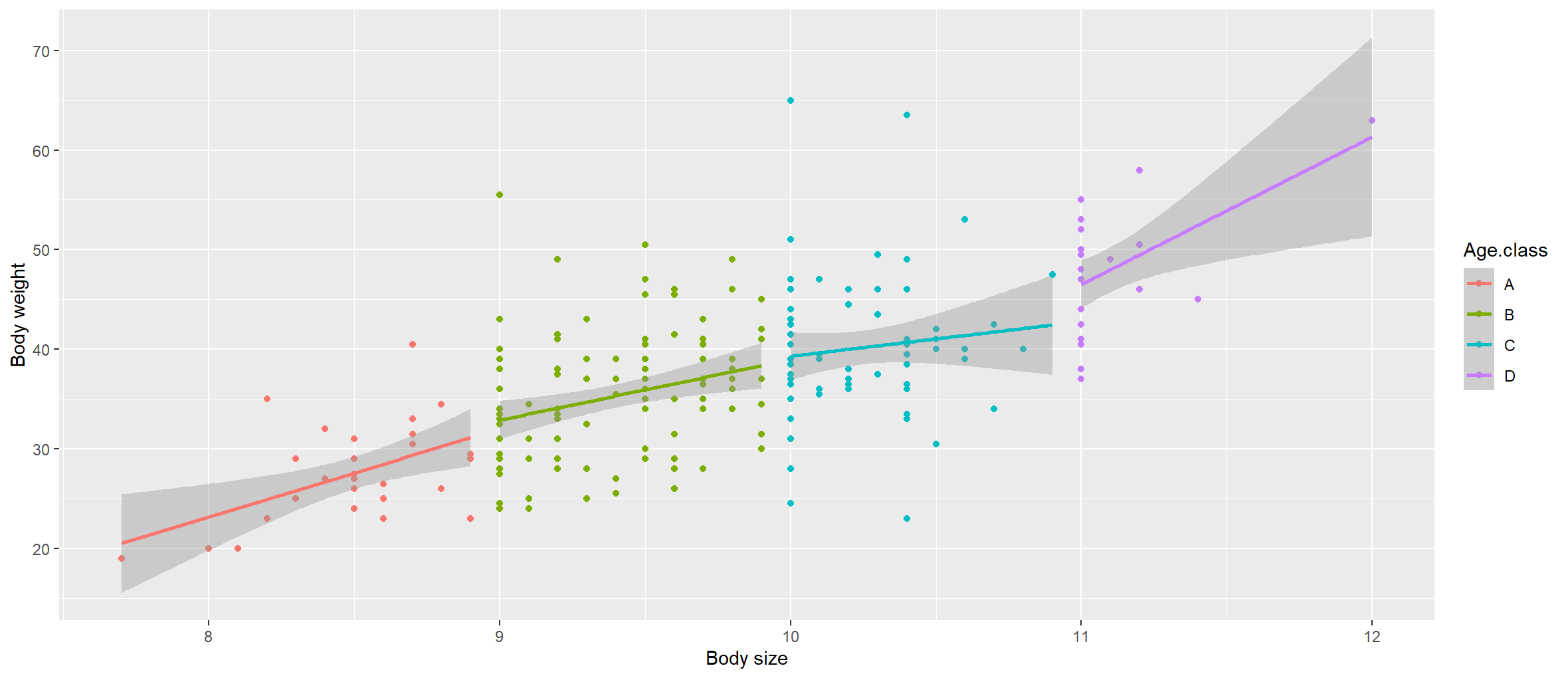

***Supplementary Figure 6:*** ***Variation of body mass with body size of adults of 4 age classes*** (Class A: [7.7:9 cm[ ; Class B : [9:10 cm[ ; Class C : [10:11 cm[; Class D [11 cm:])

**Supplementary table 11: Summary of the results of the initial Permutational Multivariate Analysis of variance model**

| Variable | Df | R² | Pseudo-F | P |
| --- | --- | --- | --- | --- |
| Session | 1 | 0.011 | 2.870 | 0.074 |
| Species | 1 | 0.042 | 11.024 | 0.020 |
| Sex | 3 | 0.010 | 2.735 | 0.087 |
| Body condition | 1 | 0.006 | 1.662 | 0.178 |
| NDVI | 1 | 0.002 | 0.556 | 0.493 |
| Age class | 3 | 0.015 | 1.272 | 0.236 |
| Breeding status | 2 | 0.029 | 3.738 | 0.020 |
| Session*Species | 1 | 0.002 | 0.565 | 0.517 |
| Session*Sex | 1 | 0.002 | 0.461 | 0.548 |
| Species*Sex | 1 | 0.016 | 4.132 | 0.025 |
| Session*Breeding status | 2 | 0.002 | 0.203 | 0.910 |
| Species*Breeding status | 2 | 0.027 | 3.558 | 0.042 |
| Session*Species*Sex | 1 | 0.005 | 1.211 | 0.271 |
| Session*Species*Breeding status | 1 | 0.009 | 2.263 | 0.181 |
| Residual | 216 | 0.824 |  |  |

**b : Alkaline phosphatase**

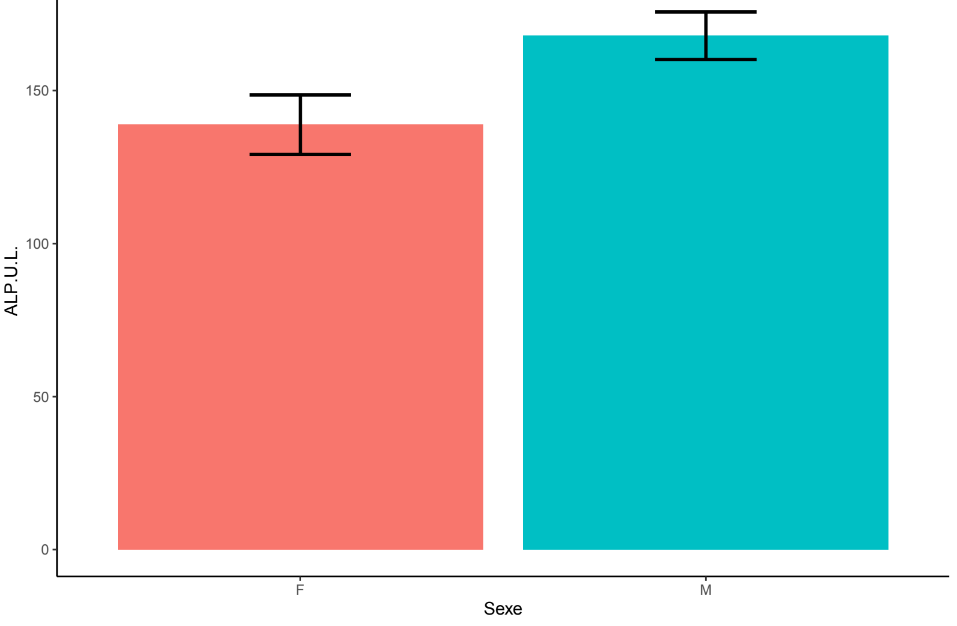

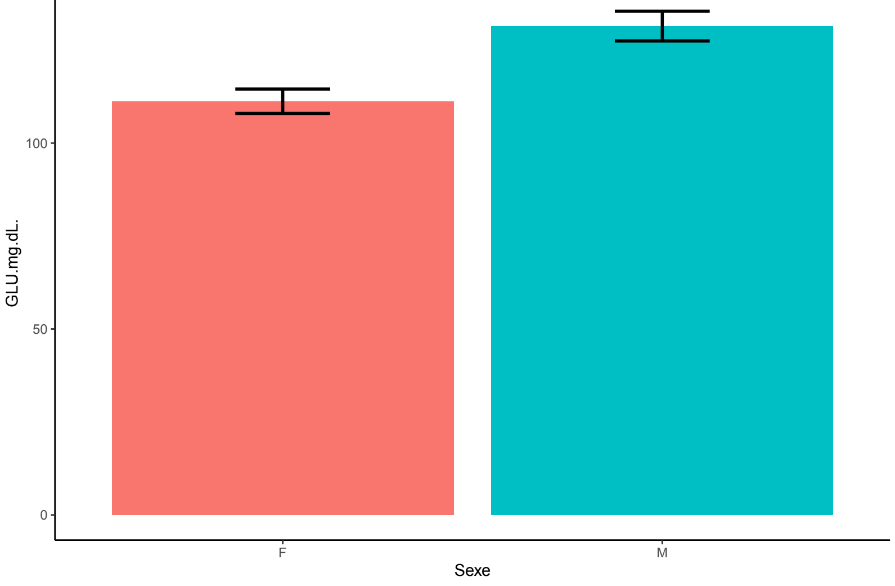

**a: Albumin**

**Supplementary Figure 7 : *Significant differences between sexes involving concentrations of Albumin and Alkaline Phosphatase (mean ± SE)***
